# Molecular Xenomonitoring (MX) allows real-time surveillance of West Nile and Usutu virus in mosquito populations

**DOI:** 10.1101/2024.04.10.588707

**Authors:** Clément Bigeard, Laura Pezzi, Raphaelle Klitting, Nazli Ayhan, Grégory L’Ambert, Nicolas Gomez, Géraldine Piorkowski, Rayane Amaral, Guillaume André Durand, Katia Ramiara, Camille Migné, Gilda Grard, Thierry Touzet, Stéphan Zientara, Rémi Charrel, Gaëlle Gonzalez, Alexandre Duvignaud, Denis Malvy, Xavier de Lamballerie, Albin Fontaine

## Abstract

**Background:** West Nile (WNV) and Usutu (USUV) virus are vector-borne flaviviruses causing neuroinvasive infections in both humans and animals. Entomological surveillance is a method of choice for identifying virus circulation ahead of the first human and animal cases, but performing molecular screening of vectors is expensive, and time-consuming.

**Methods:** We implemented the MX (Molecular Xenomonitoring) strategy for the detection of WNV and USUV circulation in mosquito populations in rural and urban areas in Nouvelle-Aquitaine region (France) between July and August 2023, using modified BG Sentinel traps. We first performed molecular screening and sequencing on excreta from trapped mosquitoes before confirming the results by detecting, sequencing and isolating viruses from individual mosquitoes.

**Findings:** We identified WNV and USUV-infected mosquitoes in 3 different areas, concurrently with the first human cases reported in the region. Trapped mosquito excreta revealed substantial virus co-circulation (75% of traps had PCR+ excreta for at least one of both viruses). *Cx. pipiens* was the most common species infected by both WNV and USUV. Genomic data from excreta and mosquitoes showed the circulation of WNV lineage 2 and USUV lineage Africa 3, both phylogenetically close to strains that circulated in Europe in recent years. Four WNV and 3 USUV strains were isolated from trapped mosquitoes.

**Interpretation:** MX strategy is easy and rapid to implement on the field, and has proven its effectiveness in detecting WNV and USUV circulation in local mosquito populations.

**Funding:** This study received funding from the Direction Générale de l’Armement (PDH 2 NBC-5-B-2212) and ARBOGEN (funded by MSDAVENIR).

**Research in context:** *Evidence before this study:* WNV and USUV circulate through complex transmission cycles involving mosquitoes as vectors, birds as amplifying hosts and several mammal species as dead-end hosts. Transmission to humans primarily occurs through mosquito bites for both viruses. Notably, WNV can also be transmitted through blood donations and organ transplants. It is estimated that a significant proportion of both WNV and USUV infections in vertebrate hosts remain unreported due to their predominantly asymptomatic nature or nonspecific clinical presentation. Nevertheless, neuroinvasive and potentially fatal disease can occur, in particular among vulnerable populations such as elderly and immunocompromised patients. In France, after its first detection in 2015, USUV has been sporadically found in eastern and southern departments, with confirmed infections in birds, mosquitoes and mammals, and few human cases described. WNV has recently caused annual outbreaks of varying intensities involving humans, equids and avifauna in French departments mainly located in the Mediterranean area. Because of low viral loads and/or brief viremia, diagnosis of both pathogens is often based on serological evidence, and few genomic data are available on strains having circulated in France. Entomological surveillance can be used as an early warning method for WNV and USUV surveillance, but is costly to implement as it requires the collection of large numbers of mosquitoes to detect virus circulation when infection rates in mosquito populations are low. Therefore, viral surveillance in France still heavily relies on human and animal surveillance, *i.e.* late indicators of viral circulation.

*Added value of this study:* This study describes the implementation of the MX (Molecular Xenomonitoring) strategy for the effective surveillance of WNV and USUV circulation within mosquito populations. MX uses of modified BG Sentinels that allow (i) trapped mosquitoes to survive for several days and (ii) corresponding mosquito excreta to be collected and preserved on filter paper. MX has demonstrated many advantages over traditional entomological surveillance. Firstly, screening excreta collectively deposited by a community of trapped mosquitoes for the presence of viruses in the first instance is time and cost efficient, as one sample is tested for viral RNA, regardless of the number and species diversity in the trap. Second, filter papers with mosquito excreta can be transported from the field to the laboratory at room temperature by regular postal mail, bringing real-time detection within reach. WNV and USUV RNA have been detected and sequenced directly from the mosquito excreta shortly after collection. Thirdly, MX adapters increase the longevity of trapped mosquitoes, thereby allowing extension of the time between trap collections and increasing the likelihood of virus shedding by infected mosquitoes. Fourthly, this approach is easy to implement in the field and requires neither a strong entomological background nor specific technical skills. All these aspects make the MX strategy a powerful, non-invasive and cost-effective tool for real-time monitoring of enzootic WNV and USUV circulation.

*Authors should describe here how their findings add value to the existing evidence.:* WNV was never detected on the Atlantic seaboard of France until October 2022. Molecular evidence of WNV circulation was obtained in 3 symptomatic horses in the Nouvelle Aquitaine region in October 2022, concomitantly with an USUV human case with no recent travel history outside the region. This was a harbinger of an increase in cases over the next year. In 2023, MX succeeded in detecting the enzootic co-circulation of WNV and USUV in rural and urban areas of Nouvelle Aquitaine, simultaneously with the first cases of WNV detected by human and animal surveillance and the first human case of USUV diagnosed in the end of July 2023. Genomic and phylogenetic information was obtained directly from trapped mosquito excreta, before information derived from animal or human surveillance. Mosquitoes from traps with PCR-positive excreta were analysed individually, which allowed to calculate infection rates in mosquitoes. WNV and USUV were isolated from single *Cx. pipiens* mosquitoes. *Cx. pipiens* was the species most commonly found positive for either viruses although WNV was also detected in *Ochlerotatus* and *Aedes* mosquitoes, including one tiger mosquito (*Ae. albopictus*) in the urban environment. We argue that the MX approach is a major asset in the early warning detection of WNV and USUV circulation to alert health policy makers and take suitable control measures.

## Introduction

West Nile (WNV) and Usutu (USUV) viruses, belong to the genus *Orthoflavivirus* (former *Flavivirus, Flavivirida*e family)^1^. They circulate in complex transmission cycles involving mosquitoes as vectors, birds as amplifying hosts, and several vertebrate species as dead-end hosts. Infections in these incidental hosts are predominantly asymptomatic or result in mild manifestations (*ie.* non-specific flu-like symptoms with a short recovery period) and therefore often go unrecognized. In a minority of cases, infections with both viruses have the potential to progress to severe and potentially fatal illnesses affecting the central nervous system (*ie*. encephalitis, meningitis), particularly in vulnerable human populations such as the elderly and immunocompromised patients. Transmission to humans occurs primarily through mosquito bites, but transmission via blood transfusion and organ transplantation from infected donors has been occasionally reported for WNV and could theoretically also occur for USUV. This makes the cryptic enzootic circulation of these viruses a sword of Damocles hanging invisibly over the heads of the most vulnerable people^2^.

Native to sub-Saharan Africa, WNVs can be classified into eight phylogenetic lineages, two of which are associated with disease in humans (1 and 2)^3^. Both recently emerged worldwide through bird migrations or their transportation by human activities. The emergence of WNV outside of Africa had a substantial impact on both public and animal health. WNV’s introduction into the United States of America (USA) in 1999 has since produced the largest outbreaks ever recorded for a neuroinvasive arboviral disease in this country, with several tens of thousands of cases and several thousand deaths^4^. WNV was introduced in Europe in the 1960s^5^. Before 2004, all WNV infections in Europe were caused by viruses from lineage 1a^6^ and were limited to sporadic cases or self-limited outbreaks. The emergence of WNV lineage 2 in central Europe in 2004 was associated with a significant increase in both the number and size of human and animal outbreaks in subsequent years^7^. In 2018, 11 countries reported a total of 1,548 local WNV infections (mostly caused by lineage 2), which exceeded the cumulative number of all infections reported between 2010 and 2017^8^. The area of known circulation of WNV lineages 1 and 2 in France has been limited to the Mediterranean basin (south-eastern part of the country), where the virus has occasionally caused symptomatic infections in humans, equids or birds since the 1960s.

The emergence of USUV in Europe was first detected in 2001 in Austria, but its presence was retrospectively documented in Italy in 1996^9^. Several African and European genotypes of USUV now circulate in Europe^10^. The emergence of USUV was first officially reported in eastern France (Haut-Rhin, Rhône and Bouches-du-Rhône departments) in 2015 by direct molecular identification in blackbirds^11^ and mosquitoes^12^, but its circulation in the country was suspected since 2009^13^. Although no human deaths have been attributed to this virus that is phylogenetically and ecologically close to WNV, USUV has caused several cases of neuroinvasive disease in Europe in recent years. USUV has been detected in blood donors, but transmission from donor to recipient has never been documented^14^.

The emergence of these two viruses always evolves towards a state of endemicity. In France, both USUV (European and African genotypes) and WNV (lineages 1 and 2) are now circulating in endemic cycles. WNV and USUV had never been detected on the Atlantic coast of France before the end of summer 2022. Serological evidence of WNV circulation was reported in Nouvelle Aquitaine with the detection of an acute infection (presence of IgM and IgG specific antibodies) in 3 symptomatic horses in October 2022, coincident with a human case of USUV with no travel history outside the region. This date marked a turning point in the epidemiology of these viruses in France and foreshadowed an increase in cases the following year.

Both the high proportion of asymptomatic infections caused by WNV and USUV and their cryptic enzootic circulation make their detection in the environment challenging. There is currently no innovative, cost-effective and easy-to-use method for the early detection of the circulation of these viruses, which is essential for triggering the systematic viral RNA screening of blood and organ donors for these viruses. Here, at the interface between entomological and environmental surveillance, we have implemented a non-invasive molecular xenomonitoring (MX) approach that uses trapped mosquito excreta to monitor the emergence and circulation of WNV and USUV in real time.

The origins of the MX approach date back to the work of Hall-Mendelin and colleagues in 2010. The authors exploited mosquito sugar feeding to detect mosquito-borne pathogens in a community of trapped mosquitoes to improve the cost/effectiveness ratio of entomological surveillance (here defined as the detection of pathogens in mosquitoes)^15^. The discovery that the excreta of infected mosquitoes contain higher virus loads than their saliva^16^, which thereby improve the sensitivity of molecular detection, led to a new arbovirus surveillance system that has proved its value in the field for several viruses^17,18^ and parasites^19^. In MX, a 3D-printed housing that fits most standard mosquito traps, provides trapped mosquitoes with a moist shelter and freely accessible sugar water and facilitates the collection of their excreta on filter paper (Supplementary file 1). Trapped mosquitoes, here used as environmental samplers, are kept alive in the field over several days, which (i) allows the time between trap collections to be extended and (ii) increases the likelihood of virus shedding by trapped infected mosquitoes. Once infected by a virus, a single mosquito can shed between 3 to 5 log10 of viral RNA per day^16^. Excreta are shipped at ambient temperature by postal mail to a laboratory. There, these samples are used to detect viral RNA in a fast, simple, efficient, and cost-effective approach. Viral genetic identities can then be rapidly revealed by sequencing viral genomic RNA contained in excreta. Importantly, the method is compatible with downstream mosquito processing for nucleic acid extraction and sequencing, as well as virus isolation allowing to obtain viral genome sequences from individual mosquitoes, to identify vector species or to estimate mosquito infection prevalence (Supplementary figure 1).

We demonstrated the value of MX in July-August 2023 in the region of Nouvelle-Aquitaine in southwest France.

## Materials and methods

### MX (Molecular Xenomonitoring) strategy

MX uses modified BG Sentinel traps (BGS, Biogents AG, Regensburg, Germany), inspired from Timmins et al.^20^ and updated from L’Ambert et al.^17^ (Supplementary figure 2). In these modified BG Sentinel traps, BGS catching bags are replaced with a 3D printed MX adapter (Supplementary file 1) attached beneath the intake funnel through a conical net and inserted into the depressurized BGS catching pipe. The MX adapter provides a safe and moisturized shelter to trapped mosquitoes with easy access to a cotton ball soaked in 10% sugar water, held in a feeder at the top inner side of the cylinder. A filter paper (Whatman, grade 1, ref. 1001-917) is placed at the bottom of the adapter to collect excreta from trapped mosquitoes.

### Study area and samples collection

The study was carried out in a ∼ 800 km² (40 x 20 km) area from either side of the Gironde estuary at the northern edge of the Bordeaux urban area, in the region of Nouvelle-Aquitaine (South-Western France) (Figure 1-A). MX traps operated on a 24 hour/7-day basis using carbon dioxide (CO_2_) as mosquito attractant. Pressurized CO_2_ bottles operated at a rate of 250 mL/min, from 8 pm to 8 am using a BG-CO_2_ timer (Biogents AG). Trapped mosquitoes were first stored at −20°C near the collection site, and then transferred to the laboratory where they were frozen at −80°C. The filter paper impregnated with mosquito excreta was removed and sent to the laboratory at room temperature by post. A new MX adapter was inserted in the trap for the following collection so that each trap was operating without discontinuity along the surveillance period. Between 20^th^ July and 3^rd^ August 2023, four MX traps (A-D) were placed in a wetland area on the right bank of the Gironde estuary. Six MX traps (E-J) were placed in a wetland area between the Dordogne and the Garonne rivers, before their confluence. From August 11^th^, the MX surveillance was extended with 3 MX traps (K-M) located in an urban area, inside the city of Bordeaux. In sites A to J, captures were conducted over 3 to 4 consecutive days, with traps collection and reconditioning for a new mosquito trapping session performed twice a week. Trap reconditioning involves collecting the adapter with live mosquitoes and replacing it with a new one. K to M traps were emptied every 2 to 4 days. An additional mosquito sampling site was implemented at a late stage on the 10^th^ of October 2023 in Châtelaillon, a town in the department of Charente-Maritime, which borders the Gironde department to the north. This sampling was carried out in the vicinity of a confirmed human case of West Nile virus.

**Figure 1:**
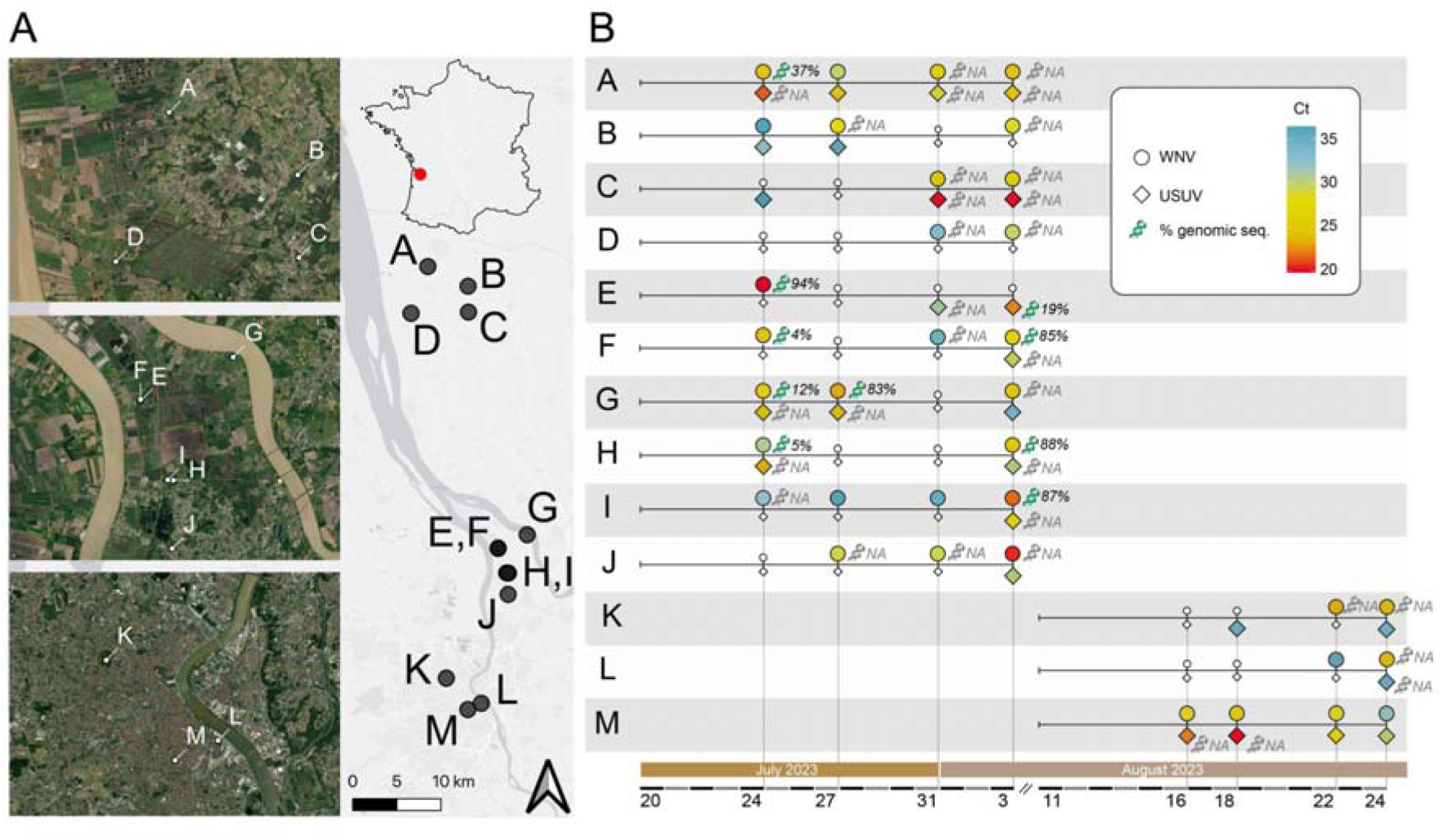
Timeline of WNV and USUV RNA detection in trapped mosquito excreta from 13 sampling sites in the department of Gironde, region of Nouvelle-Aquitaine, in the South-West of France. A) Study map with the geo-localization of the 13 sampling sites (A to M) situated on the East bank of the Gironde estuary, at the confluence of Dordogne and Garonne rivers, and within the Bordeaux agglomeration. The map was created using the free and open source QGIS geographic information system using satellite imagery from the ESRI. B) Detection of WNV (circles) and USUV (diamonds) in each sampling site (A to M) over time. Color shades correspond to cycle threshold (Ct) values, indicative of a virus load (red: low Ct and high virus load; blue: high Ct and low virus load). Little white symbols indicate no virus detection in mosquito excreta. The proportion of genomic sequence recovery from mosquito excreta at a site and collection time is represented by a DNA symbol with the genome coverage value at a sequencing depth of 50X. Grey DNA symbol indicates a failed attempt to generate sequence.

### RNA extraction from filter papers impregnated with mosquito excreta

Filter papers impregnated with mosquito excreta were stored at 4°C upon arrival to the laboratory until the RNA extraction step. Samples were not frozen and were processed in less than two weeks after reception. Briefly, each filter paper was coiled and placed at the bottom of a 14 mL plastic tubes (Falcon, ref: 352059) before being soaked in 1.5 mL of Lysis buffer RAV1 (NucleoSpin 96 virus core kit, Macherey Nagel, Düren, Germany) for 5 to 10 minutes. Ten microliters of MS2 phage were added to each tube as an internal extraction control^21^. Filter papers were then manually grinded with a 2 mL pipette until a homogeneous filter paper pulp was obtained. Several 2 mm diameter holes were then drilled in the transparent caps of the 14 mL tubes to create a colander and clipped tightly on each tube. Closed 14 mL tubes were then placed upside-down on a larger 50 mL tube (with the colander cap placed downward, at the bottom of the 50 mL tube) and centrifuged 5 min at 2,500 rpm. A 3D printed disposable spacer was used to create a space between the bottom of the 50 mL tube to avoid re-infiltration of the dry paper pulp by the flow through by capillarity after centrifugation. Flow throughs were collected and mixed with 1.5 mL 96–100% ethanol before being loaded on NucleoSpin Virus Columns in several steps. RNA extraction was then performed according to the manufacturer procedure. Eluates were stored at +4°C until used for molecular detection. This methodological step is summarized in Supplementary figure 3.

### RNA extraction from individual mosquitoes

Using RT-qPCR results on mosquito excreta, we identified traps containing mosquitoes infected with WNV and/or USUV. RNA extraction and virus detection by RT-qPCR was implemented individually for each mosquito from a selected subset of traps with positive RT-qPCR on excreta. The subset of traps were selected based on the following criteria: (*i*) the presence of a low lumber of trapped mosquitoes to reduce processing time, (*ii*) low *Ct* values obtained by RT-qPCR in excreta, indicative of a high virus load in trapped mosquitoes, (*iii*) a diversity of detection status in excreta with either the concomitant detection of both WNV and USUV, or one of these viruses alone, (iv) different location of traps (urban area, and rural area north of Bordeaux). These criteria were applied to increase the chance of virus isolation, to estimate the sensitivity and specificity of the method (working on excreta) as compared to standard entomological surveillance practices (working on mosquitoes), and to assess genomic diversity of strains circulating in different locations.

Mosquitoes were individually homogenized using 3mm tungsten beads in 400 µL of Minimum Essential Medium (MEM) supplemented with 1% penicillin–streptomycin, 1% L-glutamine, 1% Kanamycin, and 3% Amphotericin B. Homogenization was realized in a 96-well plate in a TissueLyser grinder (QIAGEN, Hilden, Germany) for 2 × 30 seconds at 30 Hz. Viral RNA was extracted with QIAamp 96 Virus QIAcube HT Kit (QIAGEN, Hilden, Germany) on a QIAcube extraction platform using 100 µL of individual mosquito lysate. RT-qPCR for WNV and USUV on individual mosquitoes was performed as described for excreta (paragraph ‘RT-qPCR on mosquito excreta’). The remaining lysate volume was stored at 4°C for subsequent viral isolation attempts to be performed for RT-qPCR-positive mosquitoes.

### RT-qPCR for for WNV and USUV

Detection of WNV and USUV genomic RNA was performed with a real-time reverse transcription polymerase chain reaction (RT-qPCR) assay. The SuperScript III Platinum One-Step qRT-PCR kit (ThermoFisher Scientific, Waltham, MA, USA) was used on a CFX96^TM^ thermal cycler, software version 3.1 (Bio-Rad Laboratories, Hercules, CA, USA). Cycling conditions were: 15 min at 50 °C, 2 min at 95 °C, 15 s at 95°C and 45 s at 60 °C (45 cycles). A 5 μL volume of RNA was added to 20 μL of mix containing 12·5 μL of 2X Reaction Mix, 0·5 μL of Superscript III RT/Platinum Taq Mix and primers and probe at the concentrations described in Supplementary table 1. A dual-target in-house assay (USUV Duo) was used to detect USUV RNA. Two RT-qPCR assays were used to detect WNV RNA. A dual-target, single dye (FAM) RT-qPCR assay (Duo WNV), combining two assays from the literature^22,23^, was used as a first-line due to its high sensitivity. Because this Duo assay cross-react with USUV, a second screening was performed with a WNV specific but less sensitive single target assay^23^ in second intention to discriminate a single WNV infection from a co-infection with both viruses when needed. A result was considered negative if the Ct value was > 40. The MS2 (internal control) RT-qPCR assay was tested on all samples.

### Viral isolation in cell culture

Remaining volume of homogenates (kept at 4°C) was retrospectively selected from mosquitoes with a RT-qPCR result lower than 38 Ct, to be inoculated on Vero African green monkey kidney cells (Vero E6 (ATCC C1008)) and C6/36 insect cells (ATCC CRL-1660). Individual mosquito’s homogenates were filtered using 0·5 mL PVDF ST ultra-free-cl millipore (Merck, Darmstadt, Germany) and diluted (1/8) in 350 µL of MEM (for Vero E6 cells) or Leibovitz’s L15 medium (for C6/36 cells) supplemented with 2.5% fetal bovine serum (FBS), 1% penicillin–streptomycin, 1% L-glutamine, 1% Kanamycin, and 3% Amphotericin B, before inoculation on confluent culture of Vero E6 and C6/36 on 6-well flat bottom cell culture plates. Individual mosquito’s homogenate inoculum was incubated 1 hour at 37°C in a 5% CO_2_ atmosphere (Vero E6 cells) or 28°C without CO_2_ (C6/36 cells) to infect cells mono-layers prior to be removed and replaced by 4 mL of MEM (for Vero E6 cells) or L-15 (for C6/36 cells) supplemented with 7% heat-inactivated FBS, 1% penicillin– streptomycin, 1% L-glutamine, 1% Kanamycin, and 3% Amphotericin B. Cell cultures were examined daily for the presence of the cytopathic effect (CPE). At post-inoculation day 5, supernatants were aliquoted and tested for the presence of WNV and USUV. Extraction and RT-qPCR were performed as described above.

### Virus sequencing

We sequenced WNV and USUV genomes from excreta and whole mosquito samples using amplicon-based approaches. We used virus-specific sets of primers to generate eight overlapping amplicons spanning the entire virus genome (Supplementary table 1, Supplementary file 3). When amplification failed for one or more amplicons, we later attempted whole genome amplification using a tiled-amplicon approach initially developed by Quick J.^24^, and Grubaugh N.D.^25^ and colleagues, with adaptations. Briefly, PrimalSeq protocol generates overlapping amplicons from 2 multiplexed PCR reactions to generate sufficient templates for subsequent high-throughput sequencing. We used primer schemes developed for WNV lineage 2^26^ and USUV^27^. For sequencing WNV and USUV genomes from virus isolates, cell culture supernatants were treated with benzonase for 1h at 37°C, extracted with no RNA carrier, and a random RT-PCR amplification was performed using the TransPlex® Complete Whole Transcriptom Amplification Kit WTA2 (Merck Millipore) following the manufacturers’ instructions. Following amplification (virus-specific or random), an equimolar pool of all amplicons was prepared for each sample, and then purified and quantified before being sonicated into 250 pb long fragments. Fragmented DNA was used for library building followed by PCR quantification. Finally, an emulsion PCR of the library pools was performed, followed by loading on 530 chips and sequencing using the S5 Ion torrent technology, following the manufacturer’s instructions (details in Supplementary file 3).

After demultiplexing, read data were analyzed with an in-house Snakemake pipeline (details in Supplementary methods). Read alignment was achieved using BWA MEM (v0.7.17) using, as a reference, the best match identified by blasting (magicblast, v1.7.7) sequencing reads using a database of flavivirus sequences including reference sequences representative of the genetic diversity of WNV (8 sequences) and USUV (13 sequences). Consensus sequences were called using a minimum coverage depth of 50x (virus-specific amplification approach) or 30x (random amplification approach).

### Phylogenetic analyses

All publicly available sequences for WNV and USUV were downloaded from the NCBI Nucleotide database, Genbank (database accessed on November 17nd, 2023). We filtered the data by: (i) excluding sequences from laboratory strains (adapted, passaged multiple times, obtained from experiments), (ii) keeping only sequences covering more than 85% of the open reading frame (ORF). The remaining sequences were trimmed to their ORF, aligned using MAFFT (version 7.511) and inspected manually using the program AliView (version 1.0). We inferred the phylogenetic relationships between public WNV and USUV genomes and the sequences generated in this study based on a Maximum-likelihood (ML) approach with IQ-Tree (version 1.6.12), using the best-fit model identified by ModelFinder and assessed branch support using an ultrafast bootstrap approximation (UFBoot2) (1000 replicates). With this first inference we identified the genotype (and phylogenetic subgroup) of all WNV and USUV sequences generated in this study. Based on these first phylogenies, we then selected smaller datasets including, for WNV, all sequences from the Central/South-West European subgroup of lineage 2 (267 sequences) and for USUV, all sequences from the Africa 3 genotype (128 sequences). Using these datasets we reconstructed time-scaled phylogenies with BEAST (v1.10.5), under the Shapiro-Rambaut-Drummond-2006 (SRD06) substitution model an uncorrelated lognormal (UCLN) clock model clock, and a bayesian skygrid coalescent models. We ran five MCMC chains of 50 million states with the BEAGLE computational library. We used Tracer (v1.7) for inspecting the convergence and mixing, discarding the first 10 % of steps as burn-in, and ensuring that estimated sampling size (ESS) values associated with estimated parameters were all >200. All xml files for these analyses are available at https://github.com/rklitting/WNV_USUV_NouvelleAquitaine_2023. Phylogenetic trees were visualized using the ggtree R package.

### Mosquito species identification through mitochondrial DNA

Morphological identification was confirmed at the species level for a subset of mosquitoes by sequencing a 710 bp region of the cytochrome oxidase subunit I (COI) using primers from Simon et al.^28^ (Supplementary table 1). Molecular identification at the species level was performed on all mosquitoes with a PCR-positive result for either WNV or USUV, and mosquitoes that failed to be identified at the species level morphologically. Five μl of mosquito excreta RNA/DNA eluates were used in a 20 μl PCR mix containing 5 μl of Hot START 5X Firepol ready-to-load DNA polymerase mix (Dutscher, Brumath, France), 2 μl of forward and reverse primers at 10 μM, and 11 μl of water. The thermal programme was: 10 min of polymerase activation at 96°C followed by 35 cycles of (i) 30 s denaturing at 96°C, (ii) 30 s annealing at 50°C and (iii) 1 min extension at 72°C, followed by a final incubation step at 72°C for 7 min to complete synthesis of all PCR products. Amplicons were subsequently sequenced using the Sanger method and the reverse primer at Microsynth AG, Lyon, France. Each sequence was visually inspected and compared with nucleotide sequences database deposited in GenBank using the BLAST algorithm (Supplementary file 4).

### Digested blood meal identification using amplicon-based metabarcoding on mosquito excreta

A ∼440 bp mitochondrial DNA section corresponding to a subfragment of COI was amplified with primers developed by Reeves et al.^29^ (Supplementary table 1) using RNA/DNA eluates extracted from trapped mosquito excreta. These primers were degenerated to selectively amplify vertebrate mitochondrial DNA while avoiding co-amplification of mosquito mitochondrial DNA. Illumina Nextera universal tails sequences were added to the 5′ end of each of these primers to facilitate library preparation by a two-step PCR approach. Six nucleotides barcodes were also inserted in the reverse primer sequences to reduce the costs by multiplexing^30^. The parameters for PCR mix and cycling were the same as for mosquito species identification described above. A 15 cycle PCR was then performed using Nextera Index Kit – PCR primers, that adds the P5 and P7 termini that bind to the dual 8 bp index tags and the flow cell. Resulting amplicons were purified with magnetic beads (SPRIselect, Beckman Coulter). Libraries were sequenced on a MiSeq run (Illumina) by Microsynth AG, Zurich, Switzerland, using MiSeq version 3 chemistry with 300 bp paired end sequencing.

The DDemux program^30^ was used for demultiplexing fastq files. Demultiplexed .fastq sequences were imported to QIIME 2 Amplicon Distribution version 2023.9 for bioinformatic analyses. The qiime2 dada2 pipeline^31^ was used for turning paired end fastq files into merged reads, filtering out Illumina adapters, denoising and removal of chimeras and filtering out replicates. Taxonomic assignment was carried out for the amplicon sequence variants (ASVs) using the qiime2 feature classifier classify consensus vsearch plugin using a database of 1,176,764 sequences gathering Fungi, Protist, and Animal COI records, recovered from the Barcode of Life Database Systems 7 March 2021 available in L’Ambert et al.^17^. Phylogenetic tree was made with the iqtree-ultrafast-bootstrap function implemented in QIIME 2 based on sequences (ASV). The script and input QIIME 2 artifacts are provided in Supplementary file 2.

All statistical analyses were performed in the statistical environment R. Figures were made using the package ggplot2 (Wickham, 2016). The Map was created using the Free and Open Source QGIS Geographic Information System using satellite imagery from the Environmental Systems Research Institute (ESRI).

## Results

### WNV and USUV were detected in trapped mosquito excreta at high rate

A total of 52 excreta samples, obtained from 13 different sites in the region of Nouvelle-Aquitaine between the end of July and August 2023, were collected and processed (Figure 1). WNV or USUV RNA was detected in 39 (75%) excreta, alone or in combination. WNV RNA was detected in 35/52 (67%) filters. USUV RNA detected in 26/52 (50%) filters. WNV and USUV RNA were both detected in the same filter in 22/52 cases (42%). At least one of the two viruses was detected at least once in all 13 sites over the study period.,. Site D (rural area in the north of Bordeaux) is the only site where WNV alone was detected. There was no site with an exclusive detection of USUV. WNV was detected at a late stage in an additional trap in Charente Maritime.

### WNV and USUV detection in trapped mosquitoes

Of the 39 filter-positive traps, we selected 7 traps (18%) to test captured mosquitoes, totalizing 364 mosquitoes. These mosquito collections originated from 6 sites at 5 different dates (site E: 25/07/2023 and 03/08/2023; sites G, I and F: 03/08/2023; site M: 18/08/2023; site K 24/08/2023). Each of the 364 mosquito was tested individually to estimate positivity rates. The positivity rate in mosquitoes from different traps ranged from 5% (N=5/94) to 18% (N=5/28) for WNV and from 2% (N=2/94) to 8% (N=5/60) for USUV (Table 1). Considering all mosquito samples, the overall positivity rate by PCR was 5% and 3% for WNV and USUV, respectively. These results may overestimate positivity rates, as we only tested mosquitoes from traps for which a PCR-positive result was obtained for WNV or USUV in the corresponding excreta. Each of the 364 mosquito was morphologically identified at the genus level: 291 (80%) and 51 (14%) were classified as *Culex* (*Cx.*) and *Aedes* (*Ae.*)/*Ochlerotatus*, respectively. The morphologically unidentified mosquitoes were subjected to molecular identification (COI sequencing). COI was also sequenced in all mosquitoes that were PCR positive for either WNV or USUV (N=17) to identify the species.

**Table 1:** Concordance of WNV and USUV detection in trapped mosquitoes and their excreta. Only mosquitoes from a subset or traps with PCR-positive results on excreta (7 collections, C1-C7) have been tested by RT-qPCR.

| Collection | Sampling date | Sites | Location | Virus detection in excreta | N mosquitoes | *Culex* | *Aedes* | Others | N (%) WNV | N (%) USUV |
| --- | --- | --- | --- | --- | --- | --- | --- | --- | --- | --- |
| C1 | 25/07/2023 | E | Estuary | WNV | 28 | 28 | 0 | 0 | 5 (18%) | 0 (0%) |
| C2 | 03/08/2023 | E | Estuary | USUV | 60 | 58 | 2 | 0 | 0 (0%) | 5 (8%) |
| C3 | 03/08/2023 | G | Estuary | WNV+USUV | 35 | 32 | 3 | 0 | 6 (17%) | 0 (0%) |
| C4 | 03/08/2023 | I | Estuary | WNV+USUV | 27 | 19 | 5 | 3 | 0 (0%) | 0 (0%) |
| C5 | 03/08/2023 | F | Estuary | WNV+USUV | 26 | 17 | 9 | 0 | 2 (8%) | 0 (0%) |
| C6 | 18/08/2023 | M | City | USUV | 94 | 83 | 4 | 7 | 0 (0%) | 4 (4%) |
| C7 | 24/08/2023 | K | City | WNV+USUV | 94 | 55 | 32 | 7 | 5 (5%) | 2 (2%) |
| **Total** |  |  |  |  | **364** | **292** | **55** | **17** | **18 (5%)** | **11 (3%)** |

*Cx. pipiens* accounted for 77% of infected mosquitoes (N=23/30): 14 were positive for WNV and 9 for USUV. Notably, no other *Culex* species were successfully identified at the molecular level among the PCR-positive mosquitoes. Among non-*Culex* mosquitoes that were found positive for either viruses, molecular identification revealed *Ochlerotatus caspius* (N=2, positive for WNV), *Ae. vexans* (N=2, positive for WNV), *Culiseta longiareolata* (N=1, positive for USUV), and *Ae. albopictus* (N=1, positive for WNV) (Supplementary file 4). The total diversity of mosquito species captured and molecularly identified in this study was limited to 5 species: *Cx. pipiens*, *Ochlerotatus caspius*, *Ae. vexans*, *Ae. albopictus* and *Culiseta longiareolata*. *Cx. pipiens* was the most frequently captured species. It was the only species –together with *Ae. albopictus*– that was trapped in every locations, both urban and rural. *Culiseta longiareolata* was the only species caught only in the urban environment.

*Cx. pipiens* showed both the highest PCR positivity rate and viral loads, independently of the virus. WNV was detected in *Cx. pipiens* with a mean Ct value of 29·8 (range 15·2-37·6 Ct), and in *Aedes/Ochlerotatus* with a mean Ct value of 33·9 (range min 33·1-34·3 Ct). USUV was detected in *Cx. pipiens* with Ct value as low as 18·8 (range 18·8-39·9 Ct, mean: 34·5 Ct), in *Culiseta longiareolata* with a Ct of 17·2 and in an undetermined *Aedes* species with a Ct of 39·3.

In collections C1, C2, C6 and C7, perfect agreement was observed between the presence of viral RNA in excreta and in the corresponding mosquitoes (Table 1). However, no USUV RNA was detected in mosquitoes from collections C3, C4 and C5, although viral RNA from both viruses had previously been detected in excreta.

A total of 34 *Culex* spp., 79 *Aedes* spp. and 7 other mosquitoes non morphologically identified at the genus level were collected in the trap from Charente Maritime in October. WNV was detected with a Ct of 35·4 in the pool of *Culex* mosquitoes (Supplementary file 2).

### High success of viral isolations from single mosquitoes with the help of data obtained from trapped mosquito excreta

All individual mosquito homogenates with a Ct < 38 were selected for attempting virus isolation. Four WNV and three USUV strains were isolated from individual mosquitoes. The success rate of virus isolation from a single mosquito was 4/18 (22%) and 3/7 (43%) for WNV and USUV, respectively; it was linked to virus load with 6/7 (86%) isolates coming from mosquitoes with a Ct<21 (Supplementary table 2). All strains were isolated on both Vero E6 and C6/36 cells, except for one USUV strain (Isolate numb. 7), which was recovered on Vero E6 cells

### Virus sequencing on excreta and mosquitoes confirmed the presence of WNV or USUV and revealed their genetic identities

To confirm the presence of WNV and USUV in virus-positive excreta and mosquitoes, we applied an amplicon sequencing approach to sequence virus genomes from these samples. For mosquito excreta, we obtained WNV and USUV genomes in 8/26 (35%) samples and 1/17 (6 %) samples, respectively. For individual mosquitoes, we obtained 4 WNV and 2 USUV genomes. We also performed sequencing on virus isolates derived from individual mosquitoes and obtained 5 WNV and 3 USUV genomes (Supplementary table 3). All WNV genomes belonged to the Lineage 2 and all USUV genomes belonged to Africa 3 genotype.

To gain insights into the geographical origin of the WNV strains circulating in Nouvelle-Aquitaine, we performed phylogenetic reconstruction using the virus genomes generated from individual mosquitoes (and from the corresponding cell culture isolate when direct sequencing from the mosquito sample failed). Within the phylogenetic tree grouping all WNV near-complete genomes (>8500 nt) available on GenBank, all sequences from Nouvelle-Aquitaine form a single (monophyletic) clade within lineage 2 (see supplementary figure 4). We obtained similar results using virus genomes obtained directly from mosquito excreta (see supplementary figure 4). In particular, our sequences belong to the Central/South-West European subgroup^32^ of that clade, which gathers sequences from Austria, Italy, Czech Republic and Germany (see Figure 2). The Nouvelle-Aquitaine clade is rooted on a sequence from Austria in 2016 and groups more broadly with strains from Austria and Slovakia from 2014-2016, the Czech Republic (2013), and recent German strains (mostly from 2018 to 2020). These results show that the strains circulating in mosquitoes in Nouvelle-Aquitaine are not directly related to L1 and L2 WNV previously identified in France in animal samples (*ie*. birds, horses).

**Figure 2:**
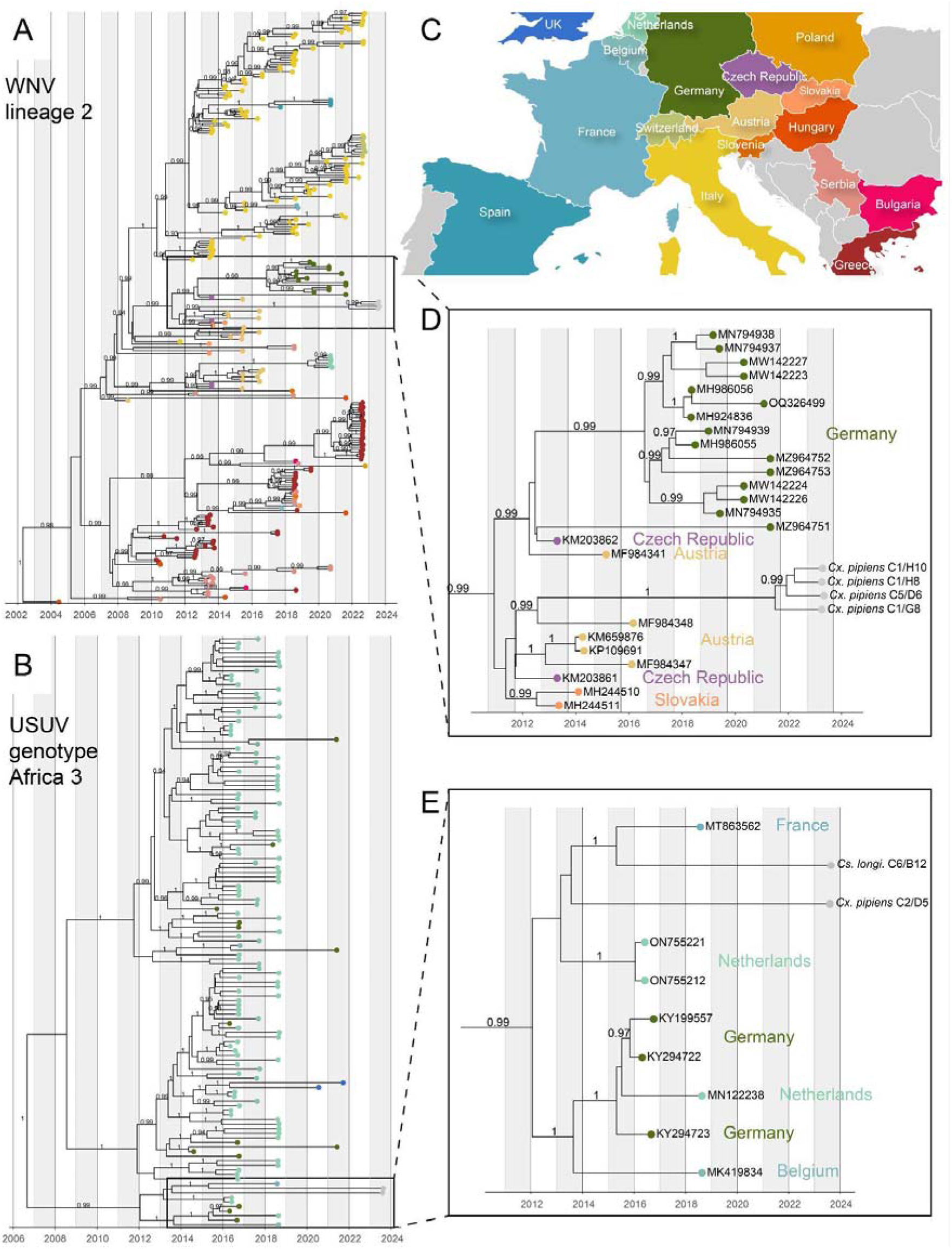
Phylogenetic relationships within WNV lineage 2 (Central European/Hungarian clade) and USUV Africa 3 genotype with a focus on sequences from Nouvelle-Aquitaine. Maximum clade credibility trees for WNV and USUV time-scaled phylogenies were reconstructed with BEAST (v1.10.5). Clade posterior supports superior to 90% are shown. A) Phylogenetic relationships within WNV from lineage 2 Central European/Hungarian clade are represented. All sequences are colored according to their geographic origin. B) Phylogenetic relationships within USUV Africa 3 genotype are represented. All sequences are colored according to their geographic origin. C) Map color highlighting the color-country correspondence used for tip coloring. D) and E) Zoom on the phylogenetic clades corresponding to WNV (D) and USUV (E) sequences from Nouvelle-Aquitaine.

Within the phylogenetic tree grouping all USUV near-complete genomes (>8500 nt) genomes available on GenBank, the virus genomes from Nouvelle-Aquitaine group together with a sequence previously sampled in Nouvelle-Aquitaine (Haute-Vienne department) in 2018, within the Africa 3 genotype (see supplementary Figure 5). We obtained similar results using virus genomes obtained directly from mosquito excreta (see supplementary figure 5). This clade from Nouvelle-Aquitaine is itself rooted by sequences sampled in Germany (2016), in Belgium (2018), and in the Netherlands (2016-2018) (Figure 2), indicating that one main lineage is currently circulating in the region.

### Sequencing digested blood from trapped mosquitoes reveals a community of vertebrates exposed to mosquitoes

DNA amplification was successful in 9 (17%) samples, all from the rural sites B (27/07/2023), D (27/07/2023), E (25 and 27/07/2023), F (27/07/2023), G (25/07/2023), H (25 and 27/07/2023) and I (25/07/2023). Sequencing generated a total of 244,424 demultiplexed read sequences across all samples with a Q20 of 92% and a median of 9,950 read sequences per sample. The total number of reads was reduced to 75 amplicons sequence variants (ASVs) with a mean length of 381 nucleotides, a mean occurrence of 1·7 ASV per sample (min: 1; first quartile: 1; third quartile: 1; max: 8) and a mean frequency of 633X (min: 2X; first quartile: 6X; third quartile: 43·5X; max: 23,879X) per sample. Following taxonomic assignment, a total of 11 ASVs (15%) were assigned to the *Arthropoda* phylum with *Chironomus riparius*, *Ochlerotatus detritus*, *Phlebotomus perniciosus*, and *Hybrizon buccatus* identified at the species level. A majority of ASVs (N=56%) were unassigned using our identity threshold, and 18 (24%) were assigned to the *Chordata* phylum. Among them 13 (17%) were identified as human DNA. As we cannot exclude that the presence of human DNA is due to contamination during sample processing, these ASVs were not considered here. Twelve ASVs were assigned to 10 vertebrate taxons: *Bos taurus* (cow), *Sus scrofa* (pig), *Canis lupus* (dog or wolf), *Felis catus* or *Felis silvestris* (cat), *Equus caballus*, *Equus ferus* or *Equus przewalskii* (horses) and *Rattus rattus* (rat) from the *Mammalia* class, *Podarcis muralis* (common wall lizard) from the *Reptilia* class, and *Streptopelia decaocto* (Eurasian collared dove), *Gallus gallus* (hen) and *Milvus migrans* or *Milvus milvus* (kites) from the *Aves* class (Figure 3). Hens, Cats, and dogs were the vertebrate species that were the most identified in both occurrence (number of study sites) and quantity (ASV counts) in these samples.

**Figure 3:**
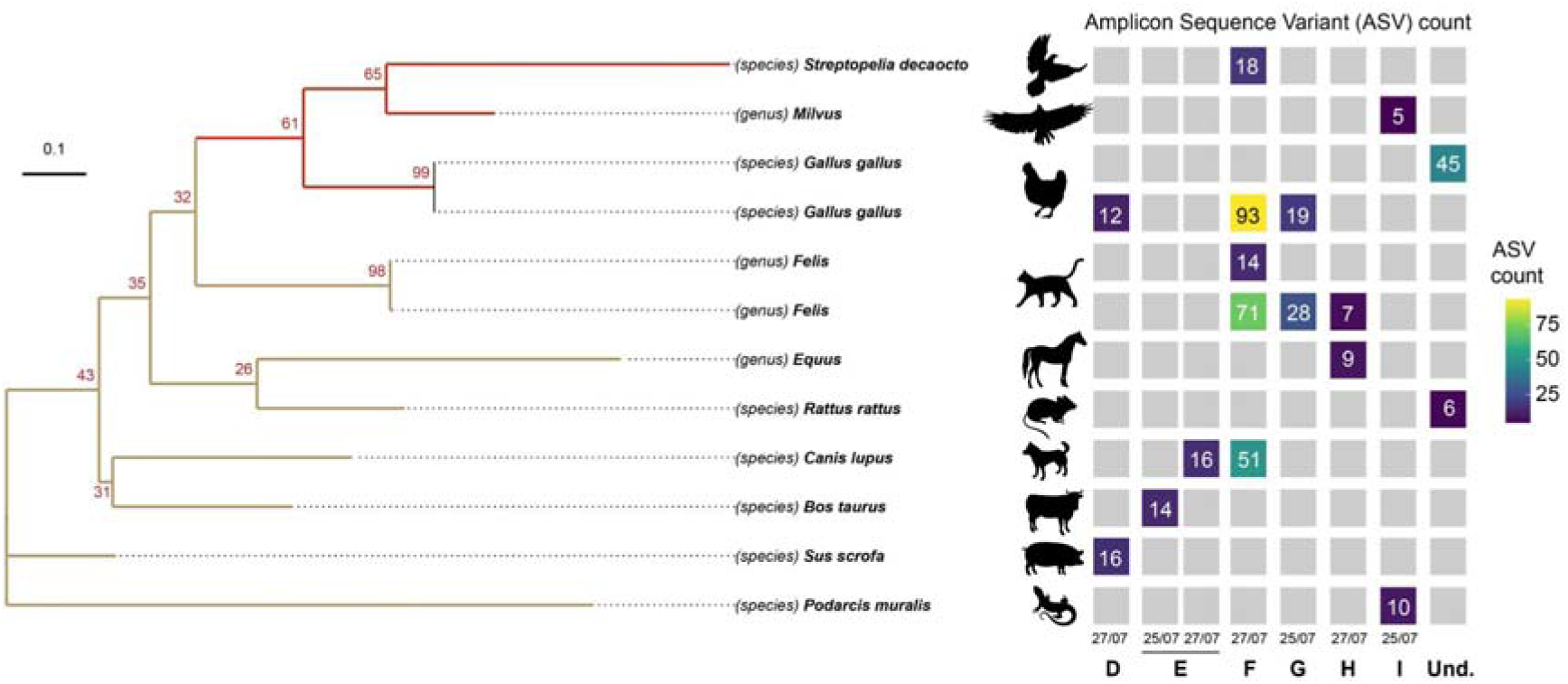
Vertebrate diversity identified based on digested blood from trapped mosquitoes. Taxons were determined at the species or genus level by comparing ASV to a sequence database using the vsearch algorithm. Genus level was chosen when an ASV matches several species inside a genus using our parameters. *Milvus*: *Milvus migrans* and *Milvus milvus*, *Felis*: *Felis catus* and *Felis silvestris*, *Equus*: *Equus caballus*, *Equus ferus* or *Equus przewalskii*. The molecular phylogenetic tree was created with the iqtree-ultrafast-bootstrap function implemented in QIIME 2 directly from ASV sequences, with 100 bootstrap replicates. This phylogenetic tree is therefore not necessarily representative of the genetic distances between these taxons. Heatmap represents the total number of ASV attributed to a taxon according to sampling sites and sampling times.

## Discussion

### A need for a new surveillance method to early detect cryptic enzootic arbovirus circulation

The accidental transmission from an organ or blood donor to recipients is one of the main risks associated with the silent circulation of enzootic neurotropic arboviruses such as WNV and, to a lesser extent, USUV. In France, systematic viral screening of all blood and organ donors is currently triggered by the diagnosis and reporting of incident WNV human cases. There is no systematic surveillance system for USUV in humans. Animal surveillance for WNV and USUV is mainly based on the reporting and diagnosis of sick horses (WNV) and dead birds (WNV, USUV). Due to the large proportion of asymptomatic infections, neither methods are fully effective in detecting the circulation of WNV/USUV in their enzootic cycles before they cause disease in humans and animals. Entomological surveillance offers non-invasive early warning capabilities, but this approach is expensive and labor intensive, requiring the processing of large numbers of mosquitoes by trained operators with strong entomological skills^33^. To our knowledge, no systematic virus screening has ever been initiated in France on the basis of entomological surveillance results.

### MX reveals the hidden circulation of enzootic arboviruses in Nouvelle-Aquitaine

Here, MX succeeded in detecting WNV and USUV enzootic transmissions in Nouvelle-Aquitaine with a high rate of detection in trapped mosquito excreta (75% of samples), one year after their suspected emergence in this area, as evidenced by serological detection in equids. In less than 2 weeks following collection, WNV and USUV genomes were detected and sequenced directly from the RNA extracted from the mosquito excreta. MX detected WNV concurrently with the confirmation of the first WNV human case in the region (end of July 2023) and a few days before the first equine case in Gironde department (4th of August). A total of 22 and 4 confirmed human cases were subsequently reported during the transmission season for WNV and USUV, respectively. MX revealed the hidden circulation of USUV concomitantly with the first confirmed human case and before the first avian case (end of July and mi-August respectively). In Italy, entomological surveillance was reported to be able to outpace the appearance of the first human infections by days to weeks^34,35^. Its early implementation in the season via MX can thus be effective in detecting enzootic arboviral circulation in endemic or emerging areas while viruses are still invisibly amplifying in animal reservoirs. The presence of WNV and USUV in Gironde, as revealed by human and animal surveillance, has made it possible to extend the detection of viral genomes in donations of human products to the surrounding Charente Maritime department from 2023 August 10^th^. This area would have not been identified as being at risk before the detection of the first WNV human case in Châtelaillon in end of August 2023. MX, which was carried out late around this human case, showed that WNV was still present in the environment two months later.

### Culex mosquitoes were major vectors in the transmission of WNV and USUV in Nouvelle-Aquitaine

*Cx. pipiens* was the most abundant species collected in both urban and rural environments and the species found with both the highest infection rate and viral loads. This species unquestionably played a leading role in the transmission of both WNV and USUV viruses in Nouvelle-Aquitaine, as it has already been reported in other transmission areas^36,37^. Here, WNV was also detected in other mosquito species. While the detection of viral RNA in mosquitoes does not necessarily confirm their infection or their ability to transmit the virus – they may simply have fed on viraemic hosts without being infected - the high USUV load recovered from a *Culiseta longiareolata* suggests a role for this ornithophilic species in the enzootic amplification cycle of these viruses. In addition, the detection of WNV in *Ae. albopictus* echoes recent findings that this urban and anthropophilic mosquito species is able to transmit WNV and USUV under experimental conditions^38^. Unlike *Culex* mosquitoes, this invasive species has been reported to have strict mammalian orientated feeding preferences^39^ and an avian component is a prerequisite for amplification of these viruses prior to infection of mosquitoes. The potential role of this species in bridging the animal reservoir to human hosts, alongside *Culex* mosquitoes, may warrant further attention if occasional blood-feeding on birds can occur.

Here, both WNV and USUV were mostly isolated from *Culex* mosquitoes exhibiting high virus loads. Infected arthropods are prime targets for virus isolation because, unlike vertebrates, they do not develop sterilizing immunity and can amplify viruses throughout their lives. Isolating viruses as they evolve and emerge worldwide can feed research activities to better understand or forecast the mechanisms that underlie their spread, pathogenicity, or adaptability in new environments.

### Easy and early access to arbovirus genomic sequences shed light on the origin and spread of viruses

Here, we obtained virus sequences directly from trapped mosquito excreta, rapidly classified the circulating WNV and USUV strains as belonging to lineage 2 and Africa 3, respectively, and confirmed these results using virus sequences obtained from single individuals. Virus sequencing from excreta samples may provide useful elements to quickly assess the potential origin of viral circulation. The resulting virus consensus genomes constitute, however, a mixing of virus populations from all virus-excreting mosquitoes –when more than one are present in the trap. In that case, analysing such genomes using phylogenetic methods may not be accurate. For that reason, we performed phylogenetic inference using virus genomes from individual mosquitoes (rather than excreta) to try to trace the origin of the virus strains identified in this study (Figure 2).

While the first detection of WNV circulation in France dates back to the 1960s^40^, limited sequence data are available to assess the spatio-temporal dynamics of circulation of the virus in the country. Here, we show that WNV sequences originating from the Nouvelle-Aquitaine region are distinct from previous L2 sequences identified in the South of France (Alpes maritimes) in 2018 from dead raptors specimens^41^ (supplementary Figure 4) and group with sequences from Austria. Based on our phylogenetic inference, the most recent common ancestor between Nouvelle-Aquitaine and Austrian sequences is approximately 10 years old (Figure 2), which makes it difficult to identify the actual timing and geographical source of the introduction of WNV into South-West France. The latter might be in Austria but may alternatively be located in another unsampled country closer to France including Italy, which has already been identified as a likely source of WNV introduction in the past^41^.

The detection of USUV in France is more recent than for WNV and dates back to 2015, when the virus was detected in the North-East (Haut-Rhin, Rhône) and South of France (Camargue), with distinct virus lineages circulating in each location, those from Rhin being apparently related to German sequences (Europe 3 genotype), those from Rhone appearing closer to sequences from Spain (Africa 2 genotype), and those from Camargue being closer to sequences from Germany and Spain (Africa 2 genotype), and the Netherlands (Africa 3 genotype). In 2018, the USUV Africa 3 genotype was identified again in Haute-Vienne, this last virus sequence is the closest phylogenetic relative of the USUV sequences identified in this study, with whom it shares a most recent common ancestor around 10 years ago (Figure 2). The long branches linking those events of virus circulation in 2018 and 2023 suggest that an important unsampled diversity of USUV genotype Africa 3 circulates in Nouvelle Aquitaine.

Altogether, our results highlight our limited knowledge of the circulating genetic diversity of WNV and USUV in France and in Europe. They call for increased genomic surveillance of arboviruses to (i) improve our understanding of the spatio-temporal circulation dynamics of these viruses at a large scale, (ii) better predict the sequential expansion of the viruses beyond the borders of Nouvelle Aquitaine and (iii) better inform public health strategies, in particular, vector management interventions.

### Molecular information contained in digested blood meals can help to reveal ecological factors involved in the emergence of these viruses

The ecological factors underlying the emergence of WNV and USUV in a Nouvelle-Aquitaine region remain unresolved. A link with the migration of birds, which are reservoirs for these viruses, can reasonably be suggested. The large fires south of Bordeaux in 2022 may also have destroyed a natural buffer zone and displaced bird populations. Mosquito excreta contains digested blood that the mosquitoes have ingested from the surrounding fauna before being caught. Sequencing regions of the selected vertebrate portion of this DNA mixture can provide insight into their local trophic preference. This method relies on catching blood engorged mosquitoes and is hampered by their rapid digestion. MX has the asset to capture vertebrate blood as it is progressively digested by trapped mosquitoes, without the need to process the mosquitoes during the digestion stage. While it cannot directly identify an animal reservoir, it does link a diversity of trapped mosquitoes to a diversity of surrounding animal hosts and, when applied at scale and combined with viral genetic information, can help to reveal the ecological forces at play in the emergence and transmission dynamics of these viruses.

### MX as a versatile early warning tool

A major drawback of entomological surveillance is that it requires time and specific knowledge. Combined with the low infection rates that typically occur in low or non-endemic areas, the method can have an unfavorable cost/effectiveness ratio that can hinder its promotion in nationwide arbovirus surveillance programs with steady funding and operational commitment. Here, we implemented a non-invasive, innovative, efficient, and cost-effective MX approach (Supplementary figure 1), at the crossroads between entomological and environmental surveillance, that succeeded in revealing the hidden circulation and the genetic identity of WNV and USUV in Nouvelle-Aquitaine. By taking advantage of excreta-based PCR testing, the MX strategy significantly accelerates the identification of potential infection hotspots, streamlining the surveillance process and facilitating more rapid and targeted public health mitigation and control measures.

## Author contributions

Clément Bigeard, Grégory L’Ambert, Guillaume André Durand, Gilda Grard, Gaëlle Gonzalez, Camille Migné, Rémi Charrel, Denis Malvy, Xavier de Lamballerie, Stéphan Zientara, Alexandre Duvignaud and Albin Fontaine designed the research. Katia Ramiara, Thierry Touzet, Grégory L’Ambert and Clément Bigeard contributed to the sample collection on the field. Laura Pezzi, Nazli Ayhan, Raphaelle Klitting, Nicolas Gomez, Géraldine Piorkowski, Rayane Amaral and Albin Fontaine performed research. Laura Pezzi, Nazli Ayhan, Raphaelle Klitting, and Albin Fontaine analyzed data. All authors participated in the redaction of the manuscript.

## Data sharing

Data are available under the NCBI BioProject number PRJNA1085973. Analysis files are available at https://github.com/rklitting/WNV_USUV_NouvelleAquitaine_2023

## Acknowledgements

We thank Thomas Canivez, Laurent Bosio, Manon Geulen, and Manon Peden from the CNR des arbovirus, Sophie Lescure, Christophe Courtin, Steeve Vernede, Hadrien Martin-Herrou from Bordeaux Métropole, Guéric Gabriel and Franck Bastit from the Communauté de Commune de Blaye and Benoît Leuret, Frédéric Jacquet and Léonard Bour from the Direction départementale de la protection des populations (DDPP) de la Gironde, Sebastien Chouin, Laurent Malnoe, Yann Renaudeau, Christian Smeraldi, and Stephane Macaud from la direction de l’Environnement et de la mobilité du département de la Charente Maritime for their help and technical assistance. The PCR tests were provided by the European virus archive-Marseille (EVAM, https://evam.european-virus-archive.com/) under the label Technological Platforms of Aix-Marseille. This work was supported by ANRS MIE, Project APP PRI 22275, Emergen-2021.

---

**Supplementary table 1: Molecular amplification systems used in this study.** Oligonucleotides sequences (primers and probes) are presented with their corresponding species, gene targets and amplicon sizes. Illumina Nextera® universal tails sequences that have been added to primers during the PCR amplification step of the arcoding method are represented in green and barcodes in blue. An adenine (A) nucleotide was added between the barcode and the primer (not mandatory).

**Supplementary table 2:** Ct values obtained from excreta, individual mosquito samples and Vero E6 and C6/36 supernatant samples obtained for isolated WNV and USUV strains. Collection/Position field correspond to the first and second column of supplementary file 4 related to virus screening in individual mosquitoes.

|  |  |  |  | Excreta |  |  | Screening<br>moustiques individual |  |  | Vero<br>E6 cells |  |  |  | C6/36 cells |  |  |
| --- | --- | --- | --- | --- | --- | --- | --- | --- | --- | --- | --- | --- | --- | --- | --- | --- |
| Isol.<br>number | Mosquito species | Virus | Collectio<br>n/Positio<br>n | USUV<br>Ct | WNV<br>Duo Ct | WNV<br>Linke<br>Ct | USUV<br>Ct | WNV<br>Duo Ct | WNV<br>Linke<br>Ct | CPE | USUV<br>Ct | WNV<br>Duo Ct | WNV<br>Linke<br>Ct | CPE | USUV<br>Ct | WNV<br>Duo Ct |
| 1 | *Cx. Pipiens* | WNV | C5/D6 | 33.2 | 26 | na | neg | 20.6 | 26.2 | + | neg | 11.4 | 13.4 | + | neg | 16.3 |
| 2 | *Cx. Pipiens* | WNV | C1/G8 | neg | 19.8 | 20.1 | neg | 16.6 | 20.1 | + | neg | 11.1 | 12.5 | + | neg | 13.9 |
| 3 | *Cx. Pipiens* | WNV | C1/H10 | neg | 19.8 | 20.1 | neg | 16.7 | 20.8 | + | neg | 11 | 12.7 | + | neg | 13.1 |
| 4 | *Cx. Pipiens* | WNV | C7/D1 | 36.9 | 24.3 | 27.5 | neg | 15.2 | 18.5 | + | neg | 11.4 | na | + | neg | 10.3 |
| 5 | *Cs. Longiareolata* | USUV | C6/B12 | 22.9 | 26.7 | neg | 17.2 | 20.4 | neg | + | 14.1 | 19.1 | neg | + | 12.5 | 21 |
| 6 | *Cx. Pipiens* | USUV | C2/A1 | 27 | neg | na | 18.8 | 20.7 | neg | + | 12.8 | 17.3 | neg | + | 12.1 | 16.2 |
| 7 | *Cx. Pipiens* | USUV | C2/D5 | 27 | neg | na | 28.6 | 31 | neg | + | 16.9 | 20.6 | neg | - | neg | neg |

**Supplementary table 3:**
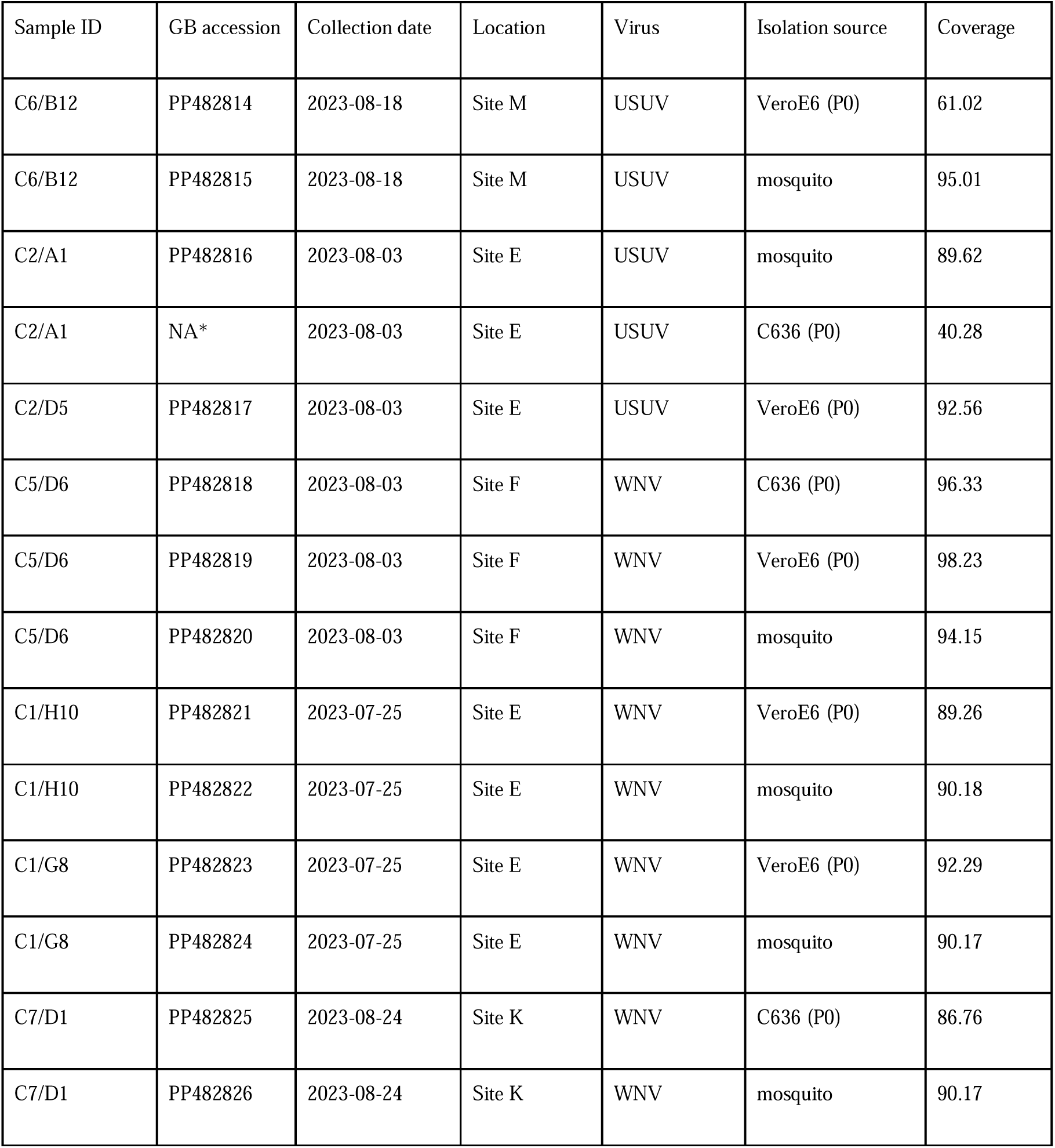
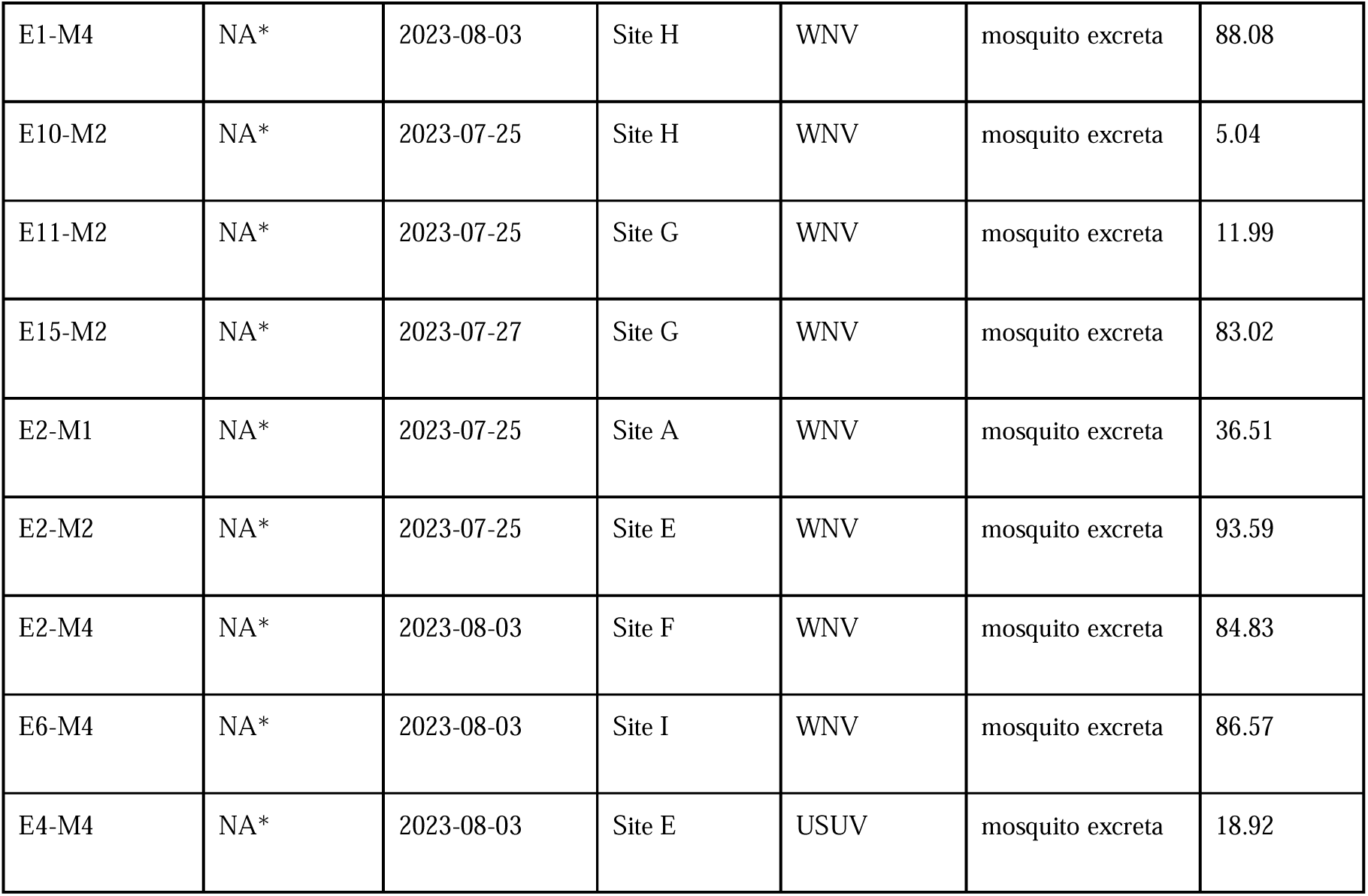
WNV and USUV sequences produced from mosquito excreta and individual mosquitoes. For each sequence, the exact source (excreta, mosquito, isolate), amplification approach, coverage at 30X (for sequences obtained from isolates) or 50X (for sequences obtained from mosquito and mosquito excreta samples), and Genbank accession number are specified. Sequencing reads for all virus genomes are available on NCBI (Bioproject ID: PRJNA1085973). Virus genomes with no Genbank accession number (NA*) are available at https://github.com/rklitting/WNV_USUV_NouvelleAquitaine_2023.

| Sample ID | GB accession | Collection date | Location | Virus | Isolation source | Coverage |
| --- | --- | --- | --- | --- | --- | --- |
| C6/B12 | PP482814 | 2023-08-18 | Site M | USUV | VeroE6 (P0) | 61.02 |
| C6/B12 | PP482815 | 2023-08-18 | Site M | USUV | mosquito | 95.01 |
| C2/A1 | PP482816 | 2023-08-03 | Site E | USUV | mosquito | 89.62 |
| C2/A1 | NA* | 2023-08-03 | Site E | USUV | C636 (P0) | 40.28 |
| C2/D5 | PP482817 | 2023-08-03 | Site E | USUV | VeroE6 (P0) | 92.56 |
| C5/D6 | PP482818 | 2023-08-03 | Site F | WNV | C636 (P0) | 96.33 |
| C5/D6 | PP482819 | 2023-08-03 | Site F | WNV | VeroE6 (P0) | 98.23 |
| C5/D6 | PP482820 | 2023-08-03 | Site F | WNV | mosquito | 94.15 |
| C1/H10 | PP482821 | 2023-07-25 | Site E | WNV | VeroE6 (P0) | 89.26 |
| C1/H10 | PP482822 | 2023-07-25 | Site E | WNV | mosquito | 90.18 |
| C1/G8 | PP482823 | 2023-07-25 | Site E | WNV | VeroE6 (P0) | 92.29 |
| C1/G8 | PP482824 | 2023-07-25 | Site E | WNV | mosquito | 90.17 |
| C7/D1 | PP482825 | 2023-08-24 | Site K | WNV | C636 (P0) | 86.76 |
| C7/D1 | PP482826 | 2023-08-24 | Site K | WNV | mosquito | 90.17 |
| E1-M4 | NA* | 2023-08-03 | Site H | WNV | mosquito excreta | 88.08 |
| E10-M2 | NA* | 2023-07-25 | Site H | WNV | mosquito excreta | 5.04 |
| E11-M2 | NA* | 2023-07-25 | Site G | WNV | mosquito excreta | 11.99 |
| E15-M2 | NA* | 2023-07-27 | Site G | WNV | mosquito excreta | 83.02 |
| E2-M1 | NA* | 2023-07-25 | Site A | WNV | mosquito excreta | 36.51 |
| E2-M2 | NA* | 2023-07-25 | Site E | WNV | mosquito excreta | 93.59 |
| E2-M4 | NA* | 2023-08-03 | Site F | WNV | mosquito excreta | 84.83 |
| E6-M4 | NA* | 2023-08-03 | Site I | WNV | mosquito excreta | 86.57 |
| E4-M4 | NA* | 2023-08-03 | Site E | USUV | mosquito excreta | 18.92 |

**Supplementary figure 1:**
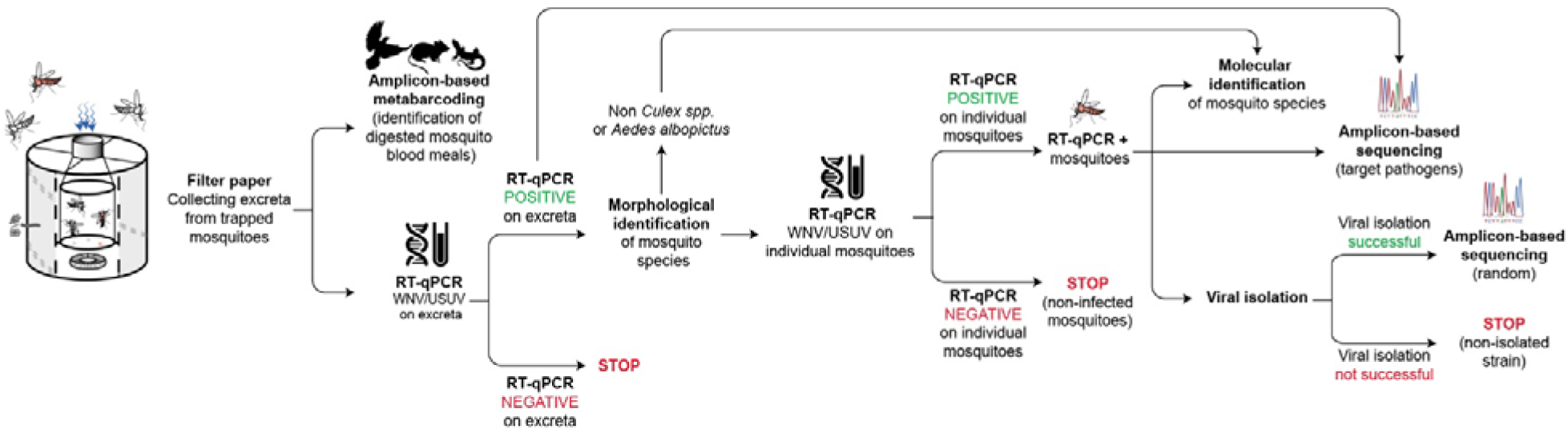
Workflow of the MX approach. Mosquitoes are captured and kept alive on the field during several days in a 3D printed shelter with a free access to sugar water. Mosquitoes are then killed and kept frozen *in situ* while the filter papers containing their excreta are sent to a laboratory at room temperature by post. Virus detection is performed at first step directly on mosquito excreta by RT-qPCR. If positive, an attempt was made to sequence the genomic RNA of the virus using amplicon-based approaches directly on the excreta. Mosquitoes from collections found positive for either viruses were then transported to the laboratory on dry ice before to be analyzed individually. Estimation of infection rates in mosquitoes, virus isolation and sequencing were performed on trapped mosquitoes on a second step.

**Supplementary figure 2:**
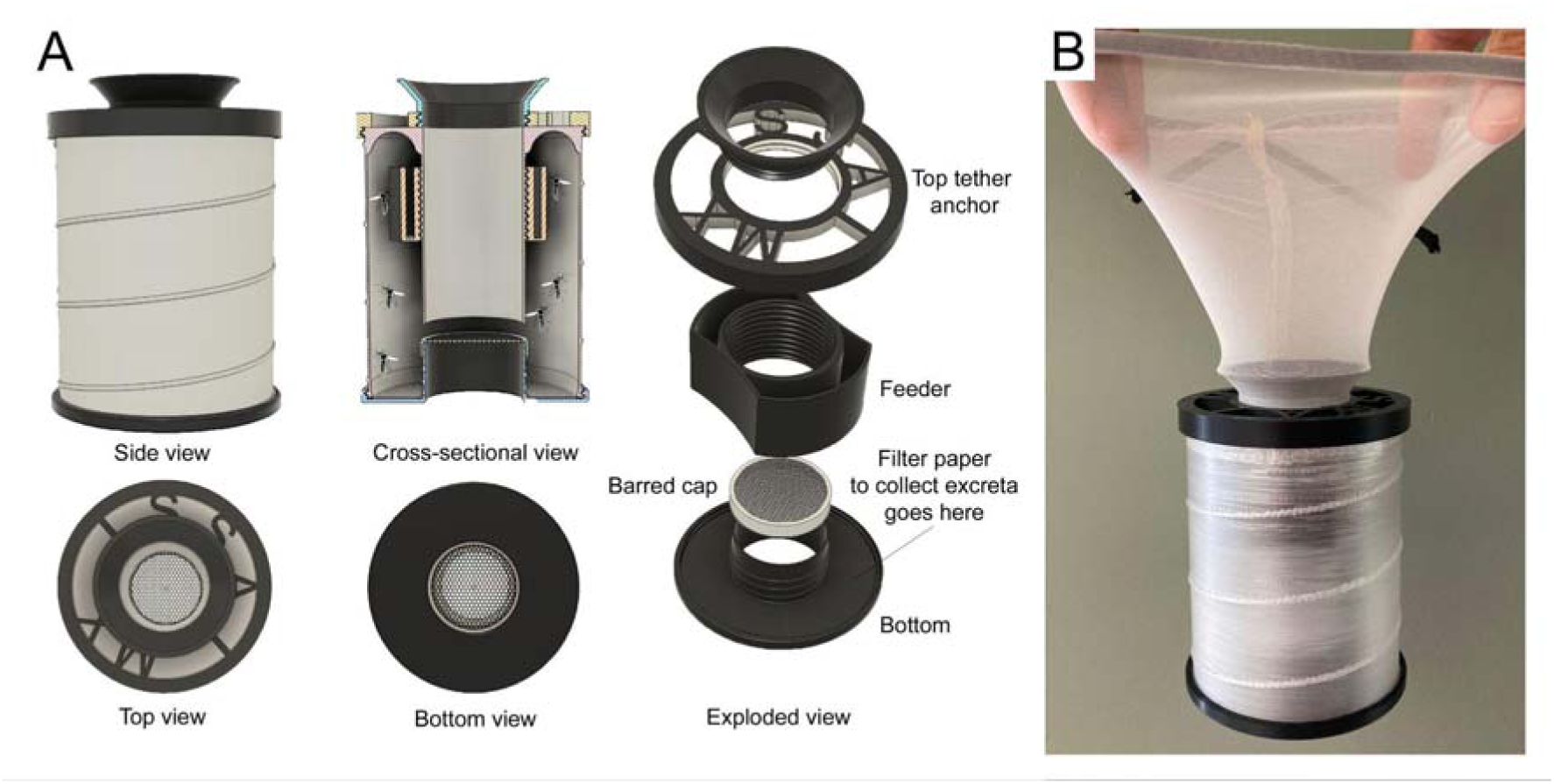
Representations of the 3D printed MX adapter designed to increase trapped mosquitoes’ longevity and to collect their excreta for an arbovirus surveillance purpose. (A) Different views of the adapter. All components are visible in the crosslJsectional and exploded views. (B) Picture of the adapter ready to be attached to the intake funnel of the BGS. The MX adapter was created on Fusion 360 (AutoDesk) and 3D printed in PLA. MX adapter 3D files (.stl format) are provided in supplementary file 1 under the Creative Commons (CC) license BY-NC-SA.

**Supplementary figure 3:**
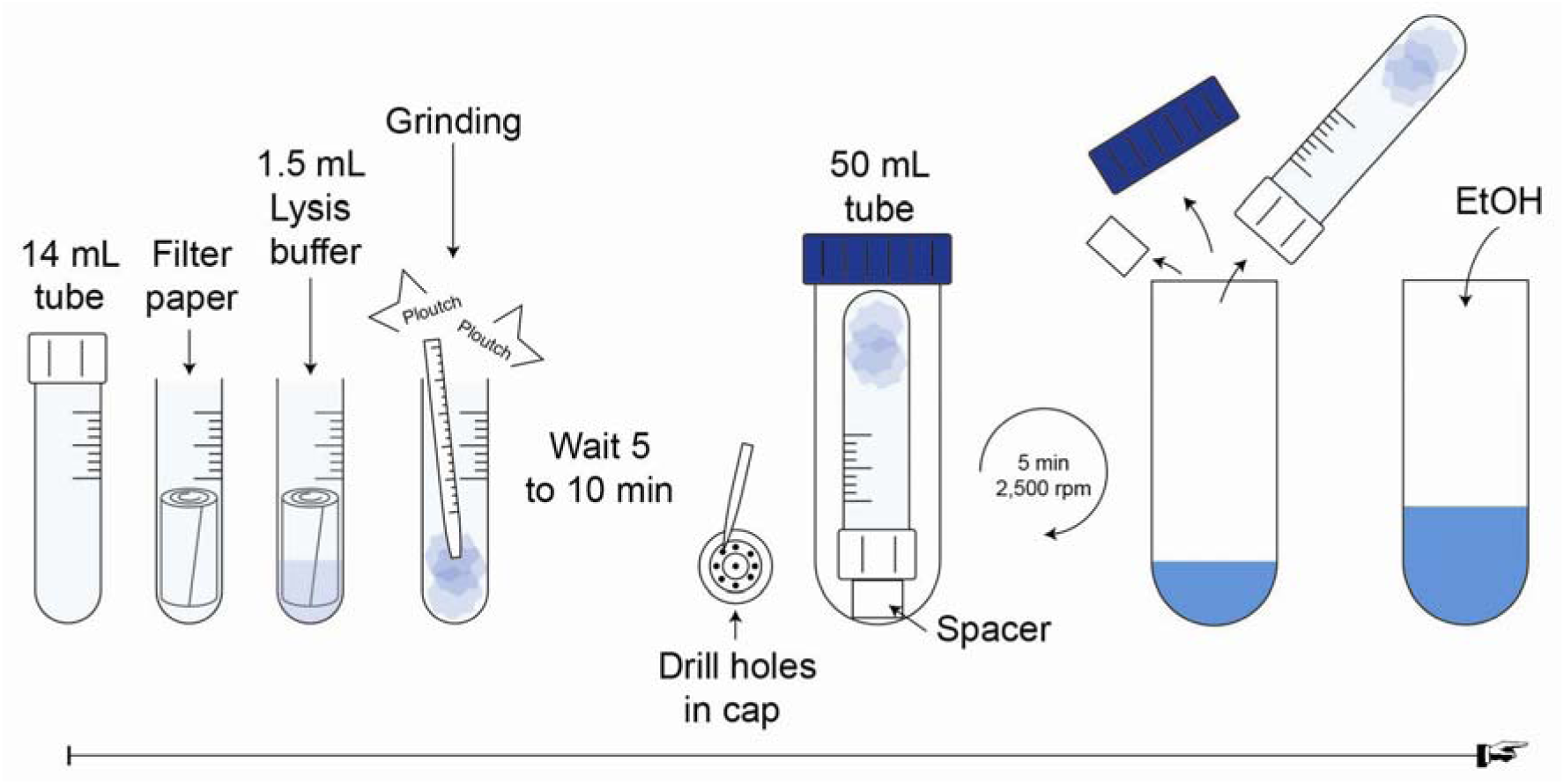
Schematic representation of the procedure to extract RNA/DNA from filter papers impregnated with mosquito excreta.

**Supplementary figure 4:**
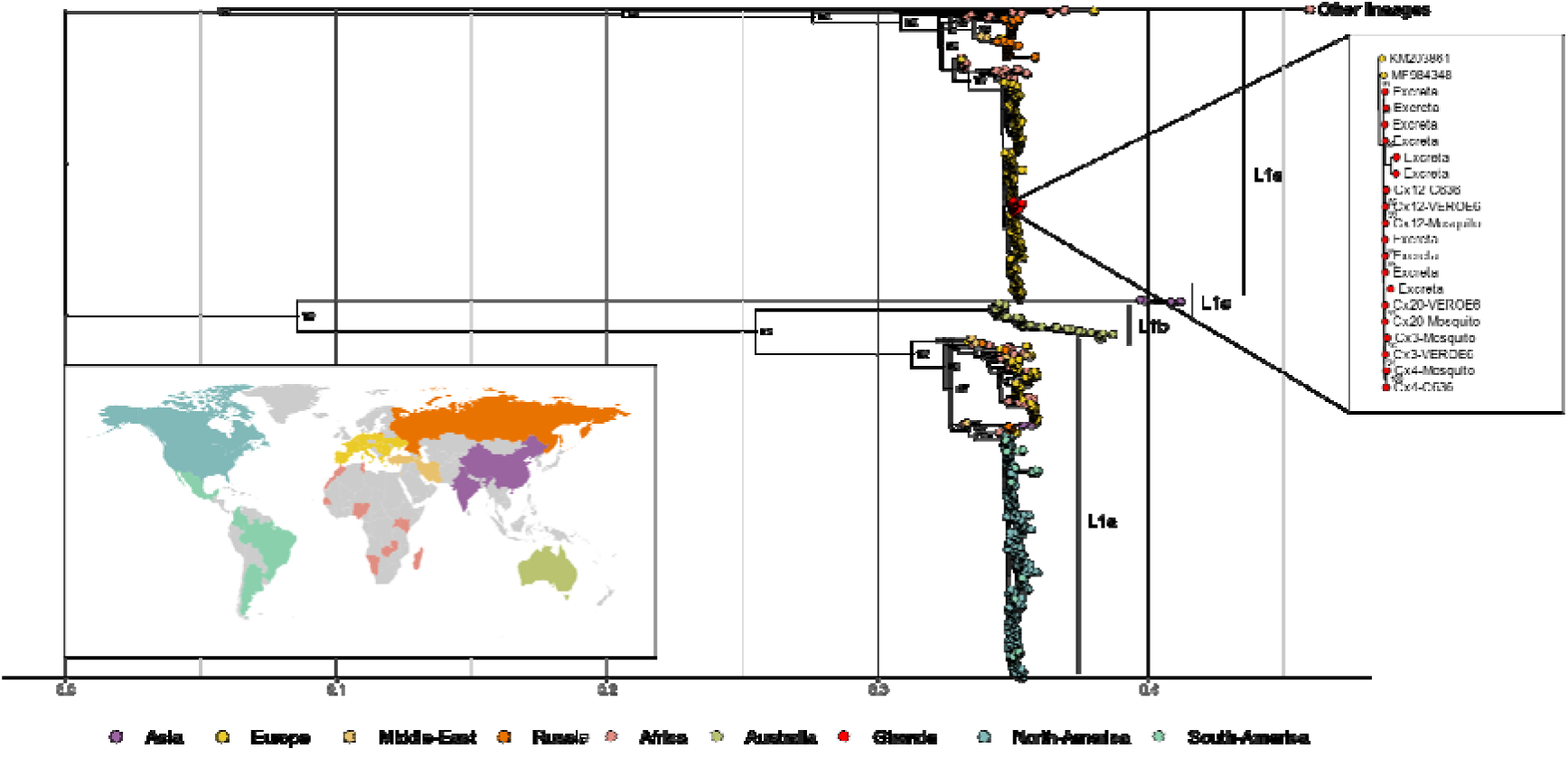
Phylogenetic relationships with WNV species with a focus on sequences from Nouvelle-Aquitaine obtained from excreta, single mosquito extract and cell culture isolates. The Maximum-likelihood phylogeny was inferred using IQ-Tree under model finder. Branch support values were calculated using UFBoot (100 replicates). Statistical supports values superior to 80% are shown for the main clades. All sequences are colored according to their geographic origin. A zoom on the clade with WNV sequences from this work is shown on the right hand side of the panel (Excreta: sequences derived from mosquito excreta, VEROE6 and C636: sequences derived from VEROE6 and C636 cell cultures, respectively, Mosquito: sequences derived from single mosquitoes).

**Supplementary figure 5:**
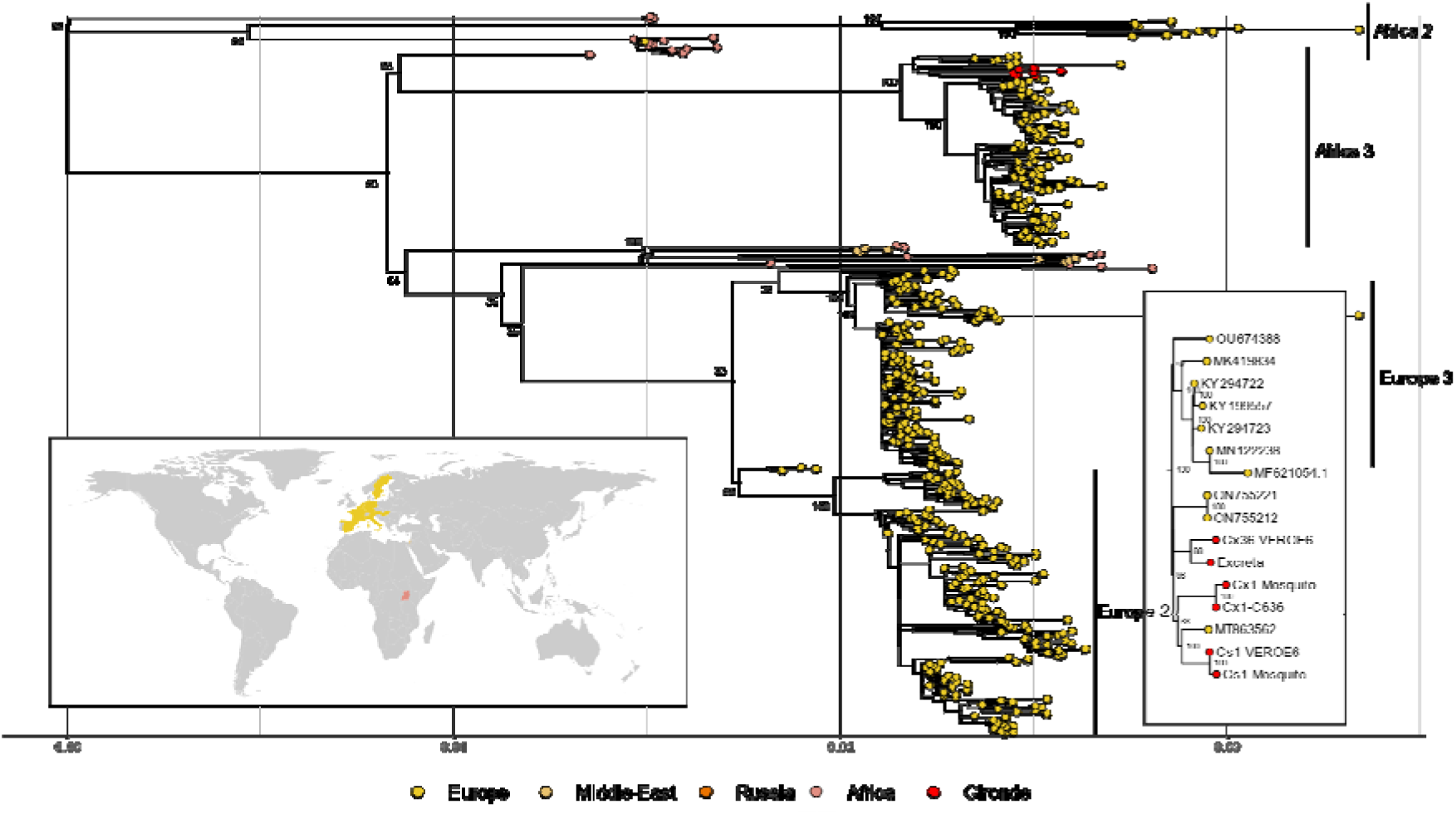
Phylogenetic relationships with USUV species with a focus on sequences from Nouvelle-Aquitaine obtained from excreta, single mosquito extract and cell culture isolates. The Maximum-likelihood phylogeny was inferred using IQ-Tree under model finder. Branch support values were calculated using UFBoot (100 replicates). Statistical supports values superior to 80% are shown for the main clades. All sequences are coloured according to their geographic origin. A zoom on the clade with WNV sequences from this work is shown on the right hand side of the panel (Excreta: sequences derived from mosquito excreta, VEROE6 and C636: sequences derived from VEROE6 and C636 cell cultures, respectively, Mosquito: sequences derived from single mosquitoes).

**Supplementary file 1: MX adapter 3D files in .stl format.** MX adapter is under the Creative Commons (CC) license BY-NC-SA (Licensees may copy, distribute, display, and make derivatives only for non-commercial purposes and by giving credits to the authors). Two versions are provided. Version 1 was used in this work. Version 2 is updated to decrease the cost and printing time.

**Supplementary file 2:** Archive comprising (i) Qiime2 code that was used in the amplicon-based metabarcoding analysis pipeline (Script_qiime2_dada2.sh), (ii) metadata file associated to the data (COI-metadata.txt), (iii) input reads (COI-paired-end.qza). All analysis and diversity metrics implemented in Qiime2 can be accessed by running Qiime2 diversity commands on data provided. All .qsv files generated by the script can be easily loaded on the Qiime2 visualizer at https://view.qiime2.org.

**Supplementary file 3: Supplementary methods.**

**Supplementary file 4: Data relative to virus detection in collections of trapped individual mosquitoes and their molecular identification at the species level.**

**Emergence timeline box:**

**1960** – Emergence of WNV lineage 1 in Europe, including in Southern France ^5^.

**1999** – Emergence of WNV lineage 1 in the United States of America. The virus has since spread southwards across the continent ^4^.

**2000** – 76 WNV cases in Equidae in Camargue, 21 deaths (Provence-Alpes-Côte d’Azur region)^42^.

**1996/2001** – Emergence of USUV in Europe ^9^.

**2001/2002** – Low level of WNV activity in Camargue as reported in sentinel birds (Provence-Alpes-Côte d’Azur region)^43^.

**2003** – 7 and 4 WNV cases in Human and Equidae, respectively, in the Var department (Provence-Alpes-Côte d’Azur region)^43^.

**2004** – Emergence of WNV lineage 2 in Europe ^7^. 37 suspected WNV cases in Equidae in Camargue (Provence-Alpes-Côte d’Azur region)^43^.

**2006** – 4 WNV cases in Equidae in Pyrénées-Orientales department (Occitanie region)^44^.

**2008** – Large WNV outbreaks in three Italian Northern Regions (Emilia Romagna, Veneto, Lombardy) with 794 cases of WNV infections in Equidae, several WNV infection detected in birds (magpies, carrion crows, and rock pigeons) and 9 WNV cases in Human^44^.

**2009/2015** – Emergence of USUV in France ^11–13^.

**2015** – 49 WNV cases in Equidae in Camargue and Hérault department (Occitanie region)^45^ and 1 WNV human case in Gard department (Occitanie region)^41^.

**2017** – 2 and 1 WNV cases in Human and Equidae, respectively, in Gard department (Occitanie region)^41^.

**2018** – High number of WNV and USUV human and animal cases in Europe. 26 and 13 WNV cases in Human, Equidae, respectively ^41^. In the avifauna: 4 WNV cases in northern goshawks, 1 in common buzzard and one in long-eared owl^41^ in Corsica and Alpes-Maritimes (Provence-Alpes-Côte d’Azur region). USUV is detected in a Lapland Owl in the Gironde (Nouvelle-Aquitaine region) department and in a blackbird in the Charente department (Nouvelle-Aquitaine region).

**2019** – 9 WNV cases in Equidae in Camargue (Provence-Alpes-Côte d’Azur region)^41^.

**2022, September** – USUV is detected in a Lapland Owl in the Dordogne department (Nouvelle-Aquitaine region).

**2022, October** – First evidence of WNV circulation on the Atlantic coast of France (3 symptomatic horses) and first human case of USUV.

**2023, July 16th** – First WNV human case in the Atlantic coast of France.

**2023, July 21th** – First USUV human case in Nouvelle-Aquitaine in 2023.

**2023, July 24th** – MX revealed the circulation of both WNV and USUV in Nouvelle-Aquitaine in 2023.

**2023, August 4th** – First WNV equine case in Nouvelle-Aquitaine in 2023.

**2023, August** – USUV is detected in a blackbird in the Charente-Maritime department (Nouvelle-Aquitaine region).

**2023, September** – USUV and WNV are detected (co-infection) in a wood pigeon in the Charente department (Nouvelle-Aquitaine region).

**2023, November** – USUV is detected in a Lapland Owl in the Dordogne department (Nouvelle-Aquitaine region).

## References

1 Postler TS, Beer M, Blitvich BJ, et al. Renaming of the genus Flavivirus to Orthoflavivirus and extension of binomial species names within the family Flaviviridae. Arch Virol 2023; 168: 224.

2 Barzon L. Ongoing and emerging arbovirus threats in Europe. J Clin Virol 2018; 107: 38–47.

3 Fall G, Di Paola N, Faye M, et al. Biological and phylogenetic characteristics of West African lineages of West Nile virus. PLoS Negl Trop Dis 2017; 11: e0006078.

4 Murray KO, Mertens E, Despres P. West Nile virus and its emergence in the United States of America. Vet Res 2010; 41: 67.

5 Hubálek Z, Halouzka J. West Nile fever--a reemerging mosquito-borne viral disease in Europe. Emerg Infect Dis 1999; 5: 643–50.

6 Koch RT, Erazo D, Folly AJ, et al. Genomic epidemiology of West Nile virus in Europe. One Health 2024; 18: 100664.

7 Bakonyi T, Ivanics E, Erdélyi K, et al. Lineage 1 and 2 strains of encephalitic West Nile virus, central Europe. Emerg Infect Dis 2006; 12: 618–23.

8 Bakonyi T, Haussig JM. West Nile virus keeps on moving up in Europe. Euro Surveill 2020; 25: 2001938.

9 Weissenböck H, Bakonyi T, Rossi G, Mani P, Nowotny N. Usutu virus, Italy, 1996. Emerg Infect Dis 2013; 19: 274–7.

10 Cadar D, Lühken R, van der Jeugd H, et al. Widespread activity of multiple lineages of Usutu virus, western Europe, 2016. Euro Surveill 2017; 22: 30452.

11 Lecollinet S, Blanchard Y, Manson C, et al. Dual Emergence of Usutu Virus in Common Blackbirds, Eastern France, 2015. Emerg Infect Dis 2016; 22: 2225.

12 Eiden M, Gil P, Ziegler U, et al. Emergence of two Usutu virus lineages in Culex pipiens mosquitoes in the Camargue, France, 2015. Infect Genet Evol 2018; 61: 151–4.

13 Vittecoq M, Lecollinet S, Jourdain E, et al. Recent circulation of West Nile virus and potentially other closely related flaviviruses in Southern France. Vector Borne Zoonotic Dis 2013; 13: 610–3.

14 Cadar D, Simonin Y. Human Usutu Virus Infections in Europe: A New Risk on Horizon? Viruses 2022; 15: 77.

15 Hall-Mendelin S, Ritchie SA, Johansen CA, et al. Exploiting mosquito sugar feeding to detect mosquito-borne pathogens. Proc Natl Acad Sci U S A 2010; 107: 11255–9.

16 Fontaine A, Jiolle D, Moltini-Conclois I, Lequime S, Lambrechts L. Excretion of dengue virus RNA by Aedes aegypti allows non-destructive monitoring of viral dissemination in individual mosquitoes. Sci Rep 2016; 6: 24885.

17 L’Ambert G, Gendrot M, Briolant S, et al. Analysis of trapped mosquito excreta as a noninvasive method to reveal biodiversity and arbovirus circulation. Mol Ecol Resour 2023; 23: 410–23.

18 Ramírez AL, Hall-Mendelin S, Doggett SL, et al. Mosquito excreta: A sample type with many potential applications for the investigation of Ross River virus and West Nile virus ecology. PLoS Negl Trop Dis 2018; 12: e0006771.

19 Minetti C, Pilotte N, Zulch M, et al. Field evaluation of DNA detection of human filarial and malaria parasites using mosquito excreta/feces. PLoS Negl Trop Dis 2020; 14: e0008175.

20 Timmins DR, Staunton KM, Meyer DB, et al. Modifying the Biogents Sentinel Trap to Increase the Longevity of Captured Aedes aegypti. J Med Entomol 2018; 55: 1638–41.

21 Ninove L, Nougairede A, Gazin C, et al. RNA and DNA bacteriophages as molecular diagnosis controls in clinical virology: a comprehensive study of more than 45,000 routine PCR tests. PLoS One 2011; 6: e16142.

22 Linke S, Ellerbrok H, Niedrig M, Nitsche A, Pauli G. Detection of West Nile virus lineages 1 and 2 by real-time PCR. J Virol Methods 2007; 146: 355–8.

23 Tang Y, Anne Hapip C, Liu B, Fang CT. Highly sensitive TaqMan RT-PCR assay for detection and quantification of both lineages of West Nile virus RNA. J Clin Virol 2006; 36: 177–82.

24 Quick J, Grubaugh ND, Pullan ST, et al. Multiplex PCR method for MinION and Illumina sequencing of Zika and other virus genomes directly from clinical samples. Nat Protoc 2017; 12: 1261–76.

25 Grubaugh ND, Gangavarapu K, Quick J, et al. An amplicon-based sequencing framework for accurately measuring intrahost virus diversity using PrimalSeq and iVar. Genome Biol 2019; 20: 8.

26 Schneider J, Bachmann F, Choi M, et al. Autochthonous West Nile virus infection in Germany: Increasing numbers and a rare encephalitis case in a kidney transplant recipient. Transbound Emerg Dis 2022; 69: 221–6.

27 Hernández-Triana LM, de Marco MF, Mansfield KL, et al. Assessment of vector competence of UK mosquitoes for Usutu virus of African origin. Parasit Vectors 2018; 11: 381.

28 Simon C, Frati F, Beckenbach A, Crespi B, Liu H, Flook P. Evolution, Weighting, and Phylogenetic Utility of Mitochondrial Gene Sequences and a Compilation of Conserved Polymerase Chain Reaction Primers. Annals of the Entomological Society of America 1994; 87: 651–701.

29 Reeves LE, Gillett-Kaufman JL, Kawahara AY, Kaufman PE. Barcoding blood meals: New vertebrate-specific primer sets for assigning taxonomic identities to host DNA from mosquito blood meals. PLoS Negl Trop Dis 2018; 12: e0006767.

30 Rašić G, Filipović I, Weeks AR, Hoffmann AA. Genome-wide SNPs lead to strong signals of geographic structure and relatedness patterns in the major arbovirus vector, Aedes aegypti. BMC Genomics 2014; 15: 275.

31 Callahan BJ, McMurdie PJ, Rosen MJ, Han AW, Johnson AJA, Holmes SP. DADA2: High-resolution sample inference from Illumina amplicon data. Nat Methods 2016; 13: 581–3.

32 Chaintoutis SC, Papa A, Pervanidou D, Dovas CI. Evolutionary dynamics of lineage 2 West Nile virus in Europe, 2004-2018: Phylogeny, selection pressure and phylogeography. Mol Phylogenet Evol 2019; 141: 106617.

33 Cristina Radaelli M, Verna F, Pautasso A, et al. Mosquito-Borne Diseases and ‘One Health’: The Northwestern Italian Experience. In: J. Rodriguez-Morales A, ed. Current Topics in Tropical Emerging Diseases and Travel Medicine. IntechOpen, 2018. DOI:10.5772/intechopen.78985.

34 Barzon L, Montarsi F, Quaranta E, et al. Early start of seasonal transmission and co-circulation of West Nile virus lineage 2 and a newly introduced lineage 1 strain, northern Italy, June 2022. Euro Surveill 2022; 27: 2200548.

35 Calzolari M, Pautasso A, Montarsi F, et al. West Nile Virus Surveillance in 2013 via Mosquito Screening in Northern Italy and the Influence of Weather on Virus Circulation. PLoS One 2015; 10: e0140915.

36 Ergunay K, Gunay F, Oter K, et al. Arboviral surveillance of field-collected mosquitoes reveals circulation of West Nile virus lineage 1 strains in Eastern Thrace, Turkey. Vector Borne Zoonotic Dis 2013; 13: 744–52.

37 Mancini G, Montarsi F, Calzolari M, et al. Mosquito species involved in the circulation of West Nile and Usutu viruses in Italy. Vet Ital 2017; 53: 97–110.

38 Martinet J-P, Bohers C, Vazeille M, et al. Assessing vector competence of mosquitoes from northeastern France to West Nile virus and Usutu virus. PLoS Negl Trop Dis 2023; 17: e0011144.

39 Faraji A, Egizi A, Fonseca DM, et al. Comparative host feeding patterns of the Asian tiger mosquito, Aedes albopictus, in urban and suburban Northeastern USA and implications for disease transmission. PLoS Negl Trop Dis 2014; 8: e3037.

40 Joubert L, Oudar J, Hannoun C, et al. [Epidemiology of the West Nile virus: study of a focus in Camargue. IV. Meningo-encephalomyelitis of the horse]. Ann Inst Pasteur (Paris) 1970; 118: 239–47.

41 Beck C, Leparc Goffart I, Franke F, et al. Contrasted Epidemiological Patterns of West Nile Virus Lineages 1 and 2 Infections in France from 2015 to 2019. Pathogens 2020; 9: 908.

42 Durand B, Chevalier V, Pouillot R, et al. West Nile virus outbreak in horses, southern France, 2000: results of a serosurvey. Emerg Infect Dis 2002; 8: 777–82.

43 Zeller H, Zientara S, Hars J, et al. West Nile outbreak in horses in Southern France: September 2004. Santé Pulibique France 2004; published online Sept 13. https://www.santepubliquefrance.fr/content/download/185268/2315428.

44 Calistri P, Giovannini A, Hubalek Z, et al. Epidemiology of west nile in europe and in the mediterranean basin. Open Virol J 2010; 4: 29–37.

45 Bahuon C, Marcillaud-Pitel C, Bournez L, et al. West Nile virus epizootics in the Camargue (France) in 2015 and reinforcement of surveillance and control networks. Rev Sci Tech 2016; 35: 811–24.

